# Structural dynamics of non-synonymous SNPs of histone methyltransferase EZH2 involved in Weaver Syndrome

**DOI:** 10.1101/2020.04.10.036517

**Authors:** Tirumalasetty Muni Chandra Babu, Vivek Keshri, Cuicui Yu, Danian Qin, Chengyang Huang

## Abstract

Enhancer of zeste homolog 2 (EZH2) is a histone H3 lysine 27 methyltransferases. Non-synonymous SNPs (nsSNPs) in the ezh2 gene may cause Weaver Syndrome that is a prominent and rare congenital disorder. Several EZH2 genetic variants have been characterized and reported, but there is no information available on their structural dynamics. Our study employs an in silico approach for structural and functional characterization of EZH2 nsSNPs. We identified 19 EZH2 nsSNPs majorly associated with Weaver Syndrome, among which four SET-domain nsSNPs including V621M, A677T, R679C, and H689Y significantly affected EZH2 structure. We conducted the triplicate of 100 ns of molecular dynamics simulations (MDS) to compare the dynamic interaction behaviors between wild-type and mutants of EZH2 with their substrate, H3K27me0 peptide. The simulation results revealed that the mutants had higher levels of structural variations as evidenced by secondary structure, density, distance plots, and principal component analysis. Compared to EZH2-WT, the mutants A677T and H689Y have shown lower binding energy with H3K27me0 peptide due to the denaturing of 310 helixes as exemplified by MM/PBSA calculations and secondary structure analysis. Our analysis shed light on mechanisms of structural variations of EZH2 nsSNPs and lay a stone to develop mutant-based therapeutic strategies for the design of target-specific scaffolds against Weaver Syndrome.

## 1. Introduction

EZH2 is a key chromatin-modifying enzyme and well-illustrated in polycomb repressive complex 2 (PRC2) involved in maintaining transcriptional repression [1]. It consists of vital subunits viz., embryonic ectoderm development (EED), retinoblastoma-binding protein 4/7 (RBBP4/7), suppressor of zeste 12 homolog (SUZ12) and enzymatic subunit enhancer of zeste homolog 2 (EZH2) [2]. Methylation of Lysine as a biological hallmark has taken place in a conserved domain of SET (Suppressor of variegation, Enhancer of Zesta trithorax), which is composed of ∼130 amino acid residues. EZH2 has two distant druggable sites on the surface of the SET-domain for binding of methyl donating cofactor, S-adenosyl methionine (SAM), and methyl-accepting substrate, H3K27me0 peptide respectively [3]. The substrate-binding site is structurally diverse since different substrates have been recognized by different histone methyltransferase (HMTase). All HMTases have shared the same binding site for co-factor, SAM although they are structurally diverse, evolutionarily conserved [4]. These vital binding pockets have become a challenge to develop target-specific drugs.

Weaver syndrome (WS) is a rare congenital disorder identified in 1974, it consists of generalized overgrowth, advanced bone age, macrocephaly, hypertelorism, variable intellectual disability and characterized facial features [5]. The patients of Weaver syndrome possibly increase susceptibility to developing cancer and particularly in early childhood a slight increase of risk has found in developing a tumor called neuroblastoma [6]. Weaver syndrome is caused by non-synonymous single nucleotide polymorphisms (nsSNP) in the ezh2 gene encoding enhancer of zeste homolog 2 (EZH2) which is a histone methyltransferase catalyzing trimethylation of histone H3 at Lysine 27 (H3K27) [7]. EZH2 activity is dependent on the proper formation of PRC2 complex with subunits like EED and SUZ12. Mutations in these subunits similarly affect EZH2-mediated pathogenicity [8,9,10]. However, more severe clinical features of WS like the development of leukemia or cerebral migration defects were significantly associated with mutations of EZH2 in the PRC2 complex [11].

Non-synonymous SNPs are coding SNPs located in the protein-coding region and alter the encoded amino acid at the mutated site which may cause structural and functional changes in the abnormal mutant protein when compared to wild-type. Totally 20604 SNPs were reported for EZH2 gene including 342 Nonsynonymous SNPs, 236 Synonymous SNPs, 189 3’-UTP SNPs, 1130 5’-UTP SNPs and 19155 Intronic SNPs. Importantly, there are 33 nsSNPs that are clinically significant, including 19 nsSNPs involved in Weaver syndrome, 3 nsSNPs involved in Malignant melanoma of the skin, 1 nsSNP involved in Lymphoma [12, 13] and 12 unspecified nsSNPs.

Because of these amino acid variations in the binding pockets, the enzyme has failed to interact with the substrate and/or cofactor in a considerable docking orientation [14, 15]. The single amino acid substitutions have been shown to alter the substrate specificity when compared to the activity of wild-type EZH2, which has a high affinity for unmethylated H3K27 and medium affinity for mono-methylated, H3K27me1 [16, 17]. Several EZH2 nsSNPs, viz., F667I, P132S, H689Y, N740K, A677T, etc., have been reported and characterized their vital role in the regulation enzyme activity [18]. However, the structural dynamics of these mutants and their interactions with H3K27me0 have not been characterized. In the present study, we perform a triplicate of 100ns MD simulations of mutants and mutant-peptide complexes to characterize the effect of amino acid substitution on the protein stability, structural variations and molecular interactions with H3K27me0 peptide. We also discuss the biochemical function and mechanism of these mutants to explain how the structural variations of EZH2 nsSNPs affect its molecular function.

## 2. Materials and Methods

### 2.1. Screening of deleterious non-synonymous SNPs in EZH2

All the reported nsSNPs for EZH2 were retrieved from dbSNP database in NCBI (http://www.ncbi.nlm.nih.gov/snp). The deleterious nsSNPs were screen out from other coding nsSNPs using two web servers, Polyphen-2 and PROVEAN. Polyphen-2 (Polymorphism Phenotyping-2) (http://genetics.bwh.harvard.edu/pph2/) [19, 20] is a pivotal web tool for the screening of deleterious nsSNPs, and it predicts the consequences of amino acid substitutions on protein structure and the corresponding function. Polyphen-2 will take protein sequence in Fasta format as input and compute the influence of amino acid variations at a given position in the query sequence. Polyphen-2 has analyzed two forms of datasets, one is HumanDiv, which compiled from all damaging allele with known function by genome-wide association studies, and another HumanVar will distinguish disease-associated mutations from all human variants. All the mutations are evaluated, qualitatively as benign, possibly or probably damaging based on the false positive rate and a threshold value of each data set [21].

PROVEAN (Protein variation effect analyzer) (http://provean.jcvi.org/index.php) [22, 23] is an online tool for the prediction of deleterious amino acid variations in the query protein sequence and the corresponding impact on the biological function of the protein. The amino acid substitutions are predicted to “deleterious” if the prediction score below the threshold value, -2.5 and “neutral” if the prediction score above the cutoff value

### 2.2. Homologous sequence identification and phylogenetic tree construction

A total of 34 sequences (33 mutants significant, and 1 wild type) were queried in NCBI non-redundant database through protein blast search analysis and top 1000 hits were considered for further filtration (>=70% query coverage and sequence identity) and analysis. The unnamed, predicted, putative, hypothetical, and partial sequences were excluded from multiple sequence alignment and phylogenetic tree. The amino acid sequences were aligned in MUSCLE program [24] and poorly aligned region were automatically treated with complete deletion method in MEGA-X software [25]. The phylogenetic tree was inferred in MEGA-X with Neighbor-Joining method, and midpoint rooted bootstrap consensus trees visualized in FigTree v1.4.4 (http://tree.bio.ed.ac.uk/software/figtree/).

### 2.3. Multiple sequence and motif analysis

To verify the conservation status of four selected nsSNP residues viz., V621, A677, R679 and H689 in the SET-domain region of various histone methyltransferases, we performed multiple sequence alignment using clustal x software and followed by motif scan using MEME suit 4.11.2 [26]. The motif analysis was carried out by selecting of maximum number of 40 with optimum motif width of 10 to 30.

### 2.4. Molecular modeling of nsSNP-based mutants and Molecular dynamics simulations

Primarily the crystal structure of the SET-domain of EZH2 (PDB id: 4mi5) [27] was retrieved from Protein data bank (http://rcsb.org) further all the water molecules were removed from the crystal structure. To investigate the structural variations among mutants (MTs) which are modeled by replacing wild-type amino acid with corresponding nsSNP residue using PyMol software and subjected to molecular dynamics simulations.

A triplicate of 100 ns time-period of molecular dynamics simulation (MDS) was performed using GROMACS 5.1.2 [28] with the OPLSA force field to understand the thermodynamic behavior of mutants and wild-type EZH2. The protein topology was created using the OPLSA program. The cubic water box was created using a simple point charge water model in the appropriated dimensions based on the surface area of protein structure [29]. The protein structure was immersed in the center of the cubic box with a minimum distance of 1.0 nm between the walls of water box and surface of any part of the protein was set up the initial process of solvation in the MD run. Further, the solvated system was neutralized with an ionic strength of 0.1 M, Na+ (sodium) and Chloride (Cl-) ions by replacing appropriated water molecules. The neutralized system was minimized through the algorithm steepest descent, minimization with a maximum number of steps 50000, long-range of electrostatic interactions was computed by particle mesh Ewald (PME). The short-range electrostatic and van der walls cut-off was set with 1.0 nm. The system was equilibrated under NVT conditions (constant number, volume, and temperature) [30] for 1000 ps at 300 K under isothermal ensemble by soft coupling with Berendsen thermostat. The internal time step was 2 femtosecond (fs) and updates the log file of every 50.0 ps. While in NPT conditions (constant number, pressure, and temperature) periodic boundary conditions were used with a constant number of particles in the systems constant pressure and the constant temperature criteria. In this simulation, the system was coupled with Parrinello-Rahmanbarostat [31] equilibrate at 1 bar pressure for 500 ps. The production dynamics for 100 ns was performed and trajectories were sampled at every 50 ps interval.

### 2.5. Residue interaction network (RIN) analysis

Residues interaction network (RIN) analysis is a handy technique, which provides in-depth information about residue and inters residue molecular interactions. RIN analysis was employed to MD simulated structures of mutant and wild-type at various time periods to characterize the molecular interactions within the binding pocket residues. Residue interaction network file of each protein was generated by using RING 2.0 webserver [35] Further, these network files were analyzed using structure-viz plug-in implemented in Cytoscape 3.7.0 software [36] to differentiates the interaction patterns of wild-type and mutant with H3K27me0 peptide.

### 2.6. Protein-peptide docking and dynamics simulations

Protein-protein docking studies were employed to enumerate the binding affinities with H3K27me0 peptide using Autodock vina program [32] with default docking parameter values. The grid box dimensions at x, y and z coordinates are prepared with 78.23, 50.14, 52.42 which are reported as binding site residues encompassing where the peptide has interactions. The docking results were ranked by binding free energy from the default score function. MD simulations were performed as described above protocol for the mutant structure complexes with H3K27me0 peptide to verify the stability of complex and dynamics of bonding interactions in comparison with the wild-type protein.

### 2.7. Analysis of Molecular dynamics trajectories

After the production dynamics, the MD trajectory files were subjected to comprehensive analysis using GROMACS built-in functions. Structural variations such as root mean square deviation (RMSD), root mean square fluctuation (RMSF), Radius of gyration (Rg) and solvent-accessible surface area (SASA) were analyzed for mutants and complexes with H3K27me0 peptide. Density plots, residue correlation distance plots and H-bond interactions between protein and peptide were computed using Gromacs functions, gmx_densmap, gmx_distmate, and gmx_hbond respectively. Secondary structural alterations were analyzed using VMD timescale plug-in [33]. All the MD plots were plotted using QtGrace software.

### 2.8. Essential dynamics

Essential dynamics or Principal component analysis (PCA) is a statistical approach to analyze the large-scale atomic motions of trajectories generated by MD simulations, which are essential for the interpretation of biological and molecular functions. Initially, the covariance matrix was generated for wild-type and mutant proteins that extract the large-scale correlated motions of protein after removing rotational and translational movements from MD trajectories. The matrix was diagonalized with an orthogonal set of eigenvectors that identifies eigenvalues. The PCA was computed with the first two principal components using Gromacs utilities [34]. The gmx_covar and gmx_anaeig, build-in functions of Gromacs were used to construct covariance matrix and principal components with trajectory files of Cα backbone atoms.

## 3. Results

### 3.1. Analysis of deleterious nsSNPs in EZH2

Totally 20604 SNPs reported on the ezh2 gene were retrieved from NCBI dbSNP. Among them, there were 1.74% nsSNPs, 1.11 % sSNP, 0.89% 3’-UTP SNPs, 5.36 % 5’-UTP SNPs, 90.90% intronic SNPs (Fig. 1A). In the nsSNPs, only 9.27% was clinically significant and the remaining 90.73 % was not shown any clinical importance (Fig. 1B). The clinically significant nsSNPs were associated more predominantly with Weaver syndrome and also involved in Malignant melanoma and Lymphoma (Fig. 1C)

The 342 retrieved nsSNPs from dbSNP were subjected to the PROVEAN web tool to evaluate the deleterious effects of specific amino acid substitutions. Our analysis shows that 109 nsSNPs have deleterious effects on EZH2 with a tolerant index score of less than -2.5 or equivalent to -2.5 and that 233 nsSNPs have neutral effects with tolerant index score more than -2.5 (Table S1).

**Figure 1.**
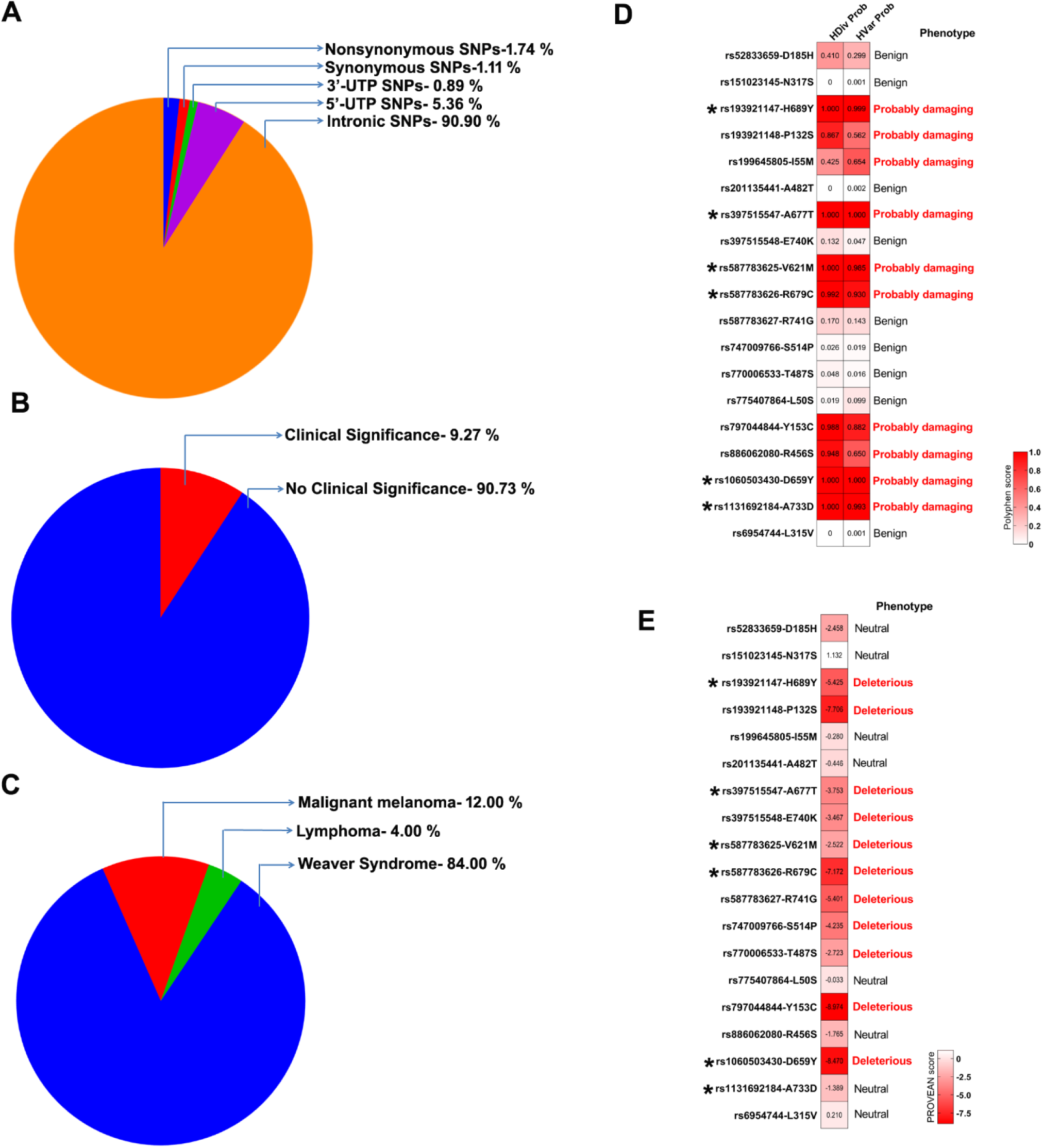
Distribution of EZH2 coding nonsynonymous SNPs (nsSNPs), coding synonymous SNPs (sSNPs), 3’-UTR SNPs, 5’-UTR SNPs and intronic SNPs **(A)**. Distribution of clinically significant and non-significant SNPs from the nsSNPs **(B)**. Distribution of disease associated nsSNPs from the clinically significant SNPs (**C**). Heatmap representation of Polyphen2 score under HDivProb and HVarProb **(D)** and PROVEAN score and their phenotype **(E)** of 19 Weaver Syndrome associated nsSNPs. The nsSNPs, localized within the SET domain region of EZH2 were highlighted with **\*** mark.

In order to verify the functional significance of amino acid replacement, these 342 nsSNPs were further submitted to Polyphen-2 with the Bayesian probabilistic models to predict whether the mutation of EZH2 is benign, possibly damaging or probably damaging. The results showed that there were 72 probably damaging nsSNPs, 50 possibly damaged nsSNPs and 124 benign nsSNPs according to HumDiv predictions (Table S2). While there were 50 probably damaging nsSNPs, 58 possibly damaging nsSNPs, and 138 benign nsSNPs according to HumVar predictions and the remaining 93 nsSNPs were not scored as any value (Table S3).

From the PROVEAN and Polyphen-2 predictions, we identified 19 Weaver syndrome-associated nsSNPs that were depicted as a heatmap in Figure 1 D and E. nsSNPs located within the SET-domain of EZH2 are functionally important therefore are major targets of an establishment of the catalytic role of corresponding amino acid replacement and effect of binding affinity with H3K27me0 peptide. There are totally 6 nsSNPs in SET-domain. We focused in particular on 4 nsSNPs including, rs587783625 (V621M), rs397515547 (A677T), rs587783626 (R679C) and rs193921147 (H689Y) that have deleterious effects on EZH2 function predicted by both of the servers and have physiological significance to Weaver syndrome. The nsSNP amino acid numbering has been used according to the amino acid sequence of reported human EZH2 (Accession number: Q15910.2).

### 3.2. Evolutionary analysis of nsSNPs in EZH2

The protein blast search analysis of EZH2 mutants detected closely related sequences from the NCBI non-redundant protein database. A total of 1000 top hits of each query were downloaded which shows a higher percentage of identity (>70%). These sequences belong to different species such as Acanthisitta chloris, Acanthochromis polyacanthus, Betta splendens, Bos taurus, Bubalus bubalis, Carlito syrichta, Danio rerio, Dasypus novemcinctus, Delphinapterus leucas, Equus caballus, Esoxlucius, Falco peregrines, Fundulu sheteroclitus, Gallus gallus, Heterocephalus glaber, Homo sapiens, Ictidomys tridecemlineatus, Kryptolebias marmoratus, Lagenorhynchu sobliquidens, Macaca nemestrina, Mus caroli, Mus musculus, Mus pahari, Notechis scutatus, Numida meleagris, Octodon degus, Odocoileus virginianus texanus, Oncorhynchus kisutch, Oncorhynchus tshawytscha, Opisthocomushoazin, Pan paniscus, Pogona vitticeps, Rattus norvegicus, Salvelinus alpines, Sarcophilus harrisii, Tupaiachinensis, Urocitellusparryii, Ursusarctos horribilis, Vulpes vulpes, Xenopus laevis, Xenopus tropicalis, Zalophus californianus, Zonotrichia albicollis and many more species. The phylogenetic tree shows the diversity of EZH2 mutant sequences and their homologs in other species (Fig. 2A). The closely related sequences are present in Castor Canadensis, Ictidomys tridecemlineatus, Urocitellus parryii, Otolemur garnettii, Pilicocolobus tephrosceles, Pan troglodytes, Papio anubis, Carlito syrichta, and Homo sapiens. Distantly related sequences are coming from Xenopus laevis, Xenopus tropicalis, Paramormyrops kingsleyae, Danio rerio, Esox lucius, Salvelinus alpines, Oncorhynchus tshawytscha etc. According to the evolutionary history this is showing that all these EZH2 sequences have been originated from a common ancestral (Fig. S1, S2) The complete list of the blast analysis is available in Table S4.

**Figure 2.**
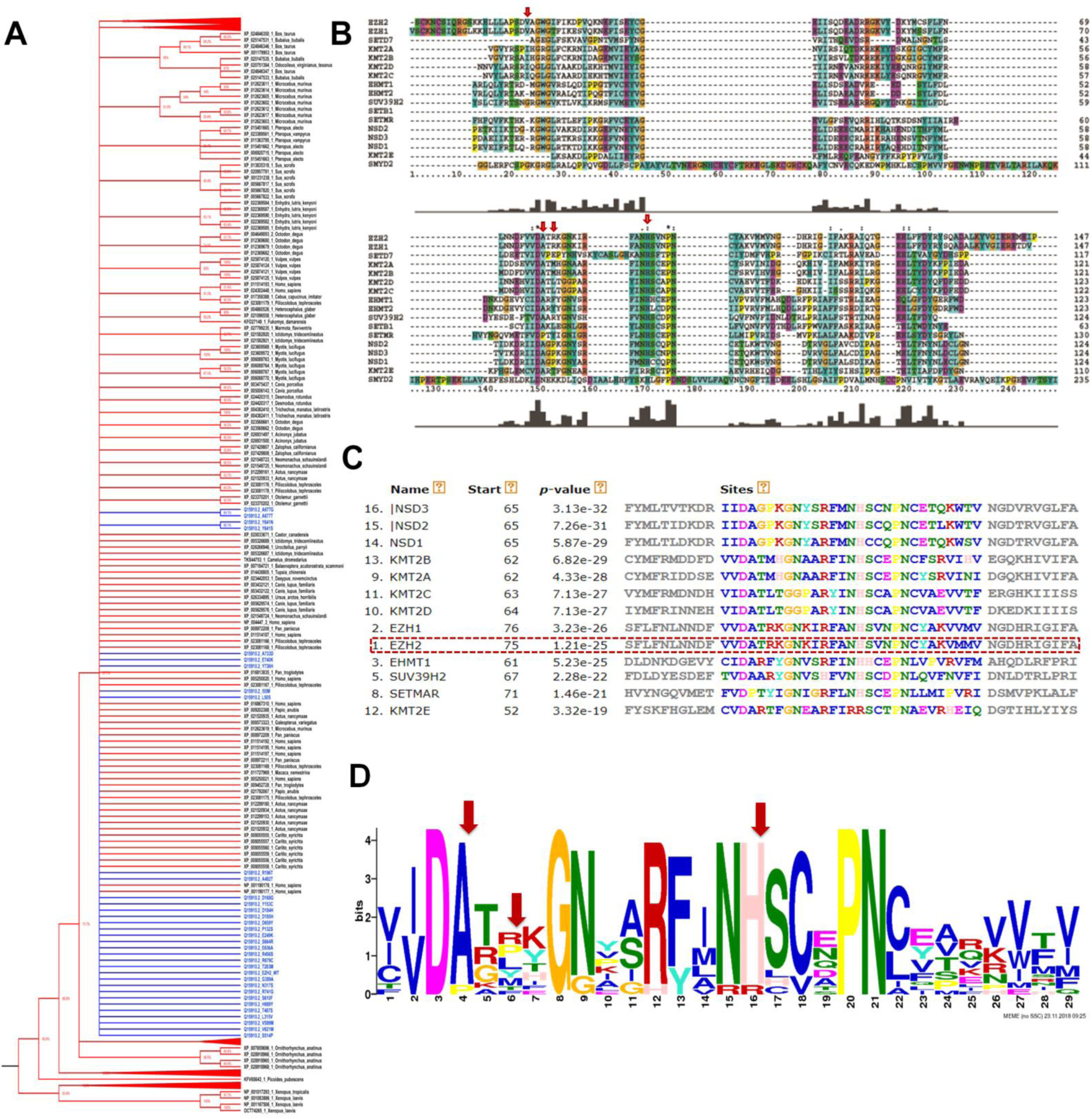
Evolutionary analyses of EZH2: The evolutionary history was inferred using the Neighbor-Joining method and bootstrap consensus tree inferred from 1000 replicates is taken to represent the evolutionary history of the taxa analyzed. The distances were computed using the p-distance method and are in the units of the number of amino acid differences per site. The rate variation among sites was modeled with a gamma distribution (shape parameter = 1). All positions containing gaps and missing data were eliminated (complete deletion). The color of the tips blue and black indicates mutant and homologous, respectively (**A**). Multiple sequence alignment of SET domain region of various histone methyltransferases including EZH2 processed through Clustal x software, the conservation of nsSNP amino acid residues were highlighted with down arrow symbol **(B)**. Motif scan analysis of SET domain region of EZH2 with the appropriated motif **(C)**, the sequential logo of the motif showing with consensus of nsSNP amino acids **(D**).

### 3.3. Multiple sequence alignment and Motif Analysis

Multiple sequence alignment of SET-domain of various HMTase including EZH2 reveals that the His 689 has shown the conservation with all HMTase except KMT2E. The Ala 677 has shown conservation with all HMTase except SETD7, SETMAR and SMYD2. The Arg 679 has shown conservation with EZH1 and SUV39H2 while Val 621 has shown conservation with only EZH1 in the corresponding positions (Fig.2B).

Commonalities in the distribution of motifs across the various histone methyltransferases were analyzed by MEME motif scan tool. It has identified six motifs across the sequence of SET-domain of EZH2. The statistical significance of motif was evaluated with their p and E values. The p value represents an estimate of how well each occurrence of matches the motif. The E value related to conservation and likewise occurrence. The results show that the nsSNP residues Ala 677, Arg 679 and His 689 has represented by motif 6 viz. VVDATRKGNKIRFANHSVNPNCYAKVMMVNG with significant and least p value of 1.21e-25 and E value of 7.2e-173 (Fig. 2C). The sequence logo was represented with highlighted nsSNP residues (Fig. 2D).

### 3.4. Molecular dynamics simulations of mutants and wild-type protein

In order to examine the structural variations, protein dynamics and stability of EZH2-WT and mutant structures i.e., V621M, A677T, R679C and H689Y, A triplicate of 100 ns time-scale of MD simulations were performed with a comprehensive analysis of dynamic trajectories after the structures were well equilibrated by NVT and NPT ensembles. The resulted MD trajectories, root mean square deviation (RMSD), root mean square fluctuation (RMSF), Radius of gyration (Rg) for backbone Cα atoms have shown wide range of variations in mutant structures when compared to wild-type EZH2.

RMSD analysis of MTs and WT structure of EZH2 with respect of initial model for the period of 50ns of MDS was plotted. From the results of the calculated RMSD plot, the WT structure showed a sharp increase of rmsd value up to ∼0.3 nm at 40 ns and equilibrated throughout the 100ns of MDS. The MT structures of A677T and H689Y have shown higher levels of structural deviations with maximum of ∼0.75 nm and equilibrated after 60 ns with minor fluctuations. While the MT structures of R679Cand V621M has shown a maximum rmsd of ∼0.5 nm and equilibrated with minor fluctuations (Fig.3A).

To enumerate the mutational effect on the flexibility of backbone of Cα atoms the root mean square fluctuations were analyzed for WT and MTs. The comparative analysis of RMSF profile indicated that shown higher fluctuations in MT structure when compared to WT structure. From the RMSF plot, the WT has shown the fluctuations throughout the enzyme sequence within the range of 0-0.35 nm, along with, the maximum RMSF value-0.35 nm in the amino acid regions of 525-540, 620-625 and 660-670. Whereas MT structures of H689Y, V621M, R679C and A677T have shown maximum fluctuations with 0.65 nm, 0.65nm, 0.55nm and 0.50 nm at the amino acid regions of 735-740, 530-540, 530-535 and 530-542, respectively (Fig. 3A).

**Figure 3.**
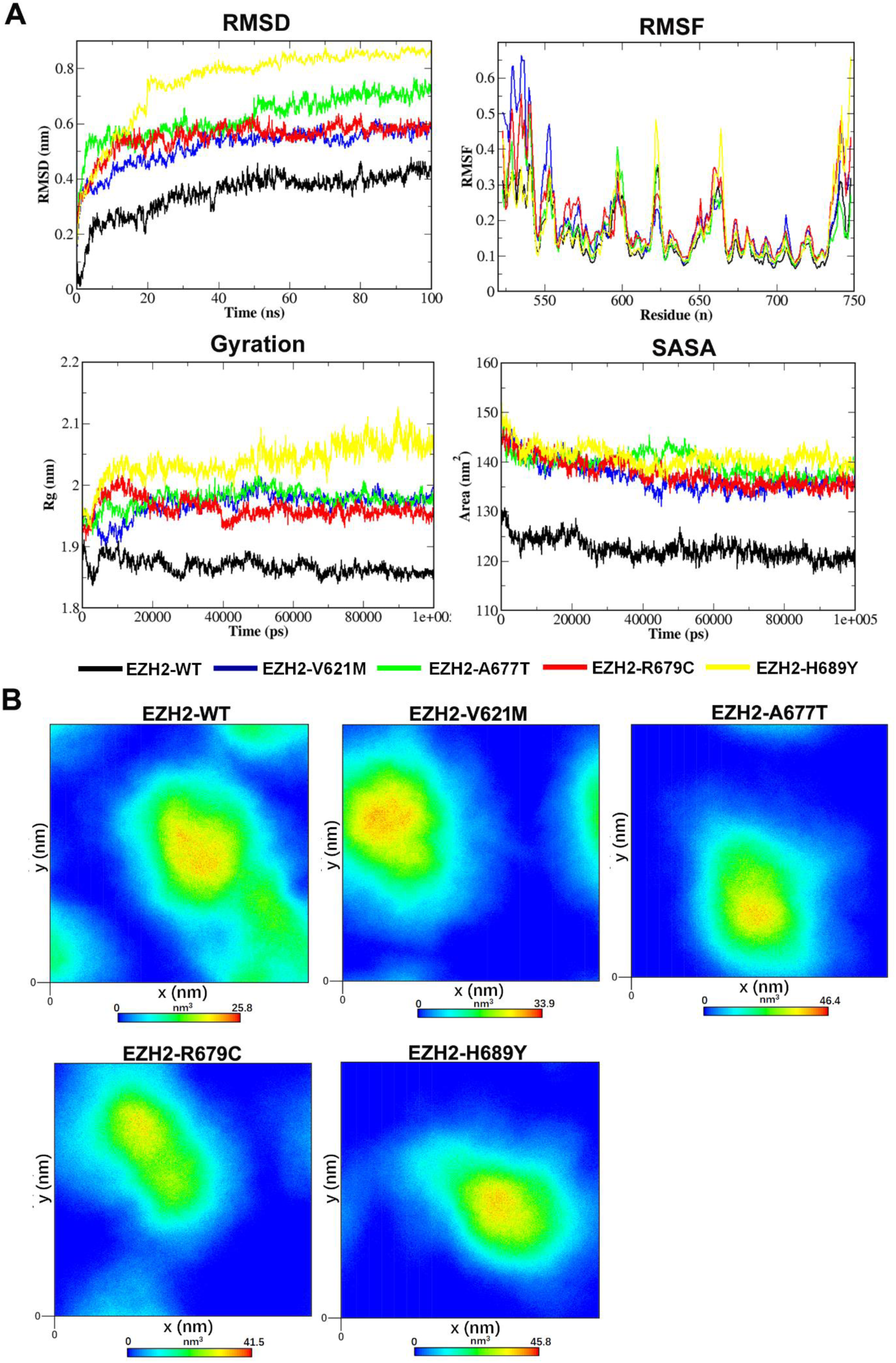
The backbone Cα atoms root mean square deviation (RMSD) values, root mean square fluctuation of backbone Cα atoms (RMSF) radius of gyration (Rg) of backbone atoms and solvent accessible surface area (SASA) of EZH2-WT, EZH2-V621M, EZH2-A677T, EZH2-R679C and EZH2-H689Y during the 100 ns molecular dynamics simulation time period **(A)**. Density plot EZH2-WT and mutant structure V621M, A677T, R679C and H689Y during 100 ns of MD simulations **(B)**.

Radius of gyration (Rg) is a parameter which defines the structural compactness with protein folding and unfolding processes [37]. In the present analysis the computed Rg values of WT and MT structures were plotted and depicted in Figure 3A. The Rg plot demonstrates that the MT structure H689Y have shown higher variations with maximum Rg value with in the range of 2.0-2.1 nm. The MT structures of V621M, A677T and R679C. have shown Rg values within the range of 1.90-2.00 nm when compared to WT structure, which shows the Rg value in the range of 1.85-1.90 nm throughout the 100ns of MDS. Therefore, the Rg results evidenced that mutational residue induces the protein unfolding that led to increase of Rg values in MT structures when compared to WT structure.

Solvent Accessible surface area (SASA) is a geometric measure of protein surface exposed to the solvent environment [38]. The SASA of MTs and WT were analyzed during 100ns of MDS. The results showed that the MT structures of V621M, A677T, R679C and H689Y have higher exposure of solvent with an average area of 140 nm2 when compared to WT structure, which shows an average area of 125 nm2 (Fig.3A).

Furthermore, the atomic density distributions were plotted with x and y axis of density values (nm3) based on the default SPC water box density to verify the changes in density profile and orientations of MTs. As shown in Figure 3B, MTs showed the density values in the order of A677T<H689Y<R679C<V621M that are significantly higher than the density value of wild type (25.8 nm3). It is evidenced that EZH2 MTs were structurally deleterious when compared to wild type (Fig.3B)

The perturbations of secondary structural patterns of EZH2-WT and its mutants were analyzed using VMD time-scale plug-in tool. The analysis report demonstrates the fluctuation of secondary structure elements throughout the 100ns of MD simulations. In the structure of EZH2-WT, turns were observed in the region of 526-530 amino acid residues up to 10 ns, after a conformational drift of turns to extended conformer up to 30 ns, further the partial replacement of extended conformer to turns in the same region throughout the 50ns of MDS, which can be consider has one significant secondary structure fluctuation. The 310 helixes were found in the region of 595-600 amino acid residues up to 10 ns further, a conformational replacement of 310 helixes to α helixes towards 50ns of MDS. These can be two significant secondary structure fluctuations and depicted as change 1 and change 2 at 5ns, 20ns, 40 ns and 50ns time-scale points (Fig. 4A).

**Figure 4.**
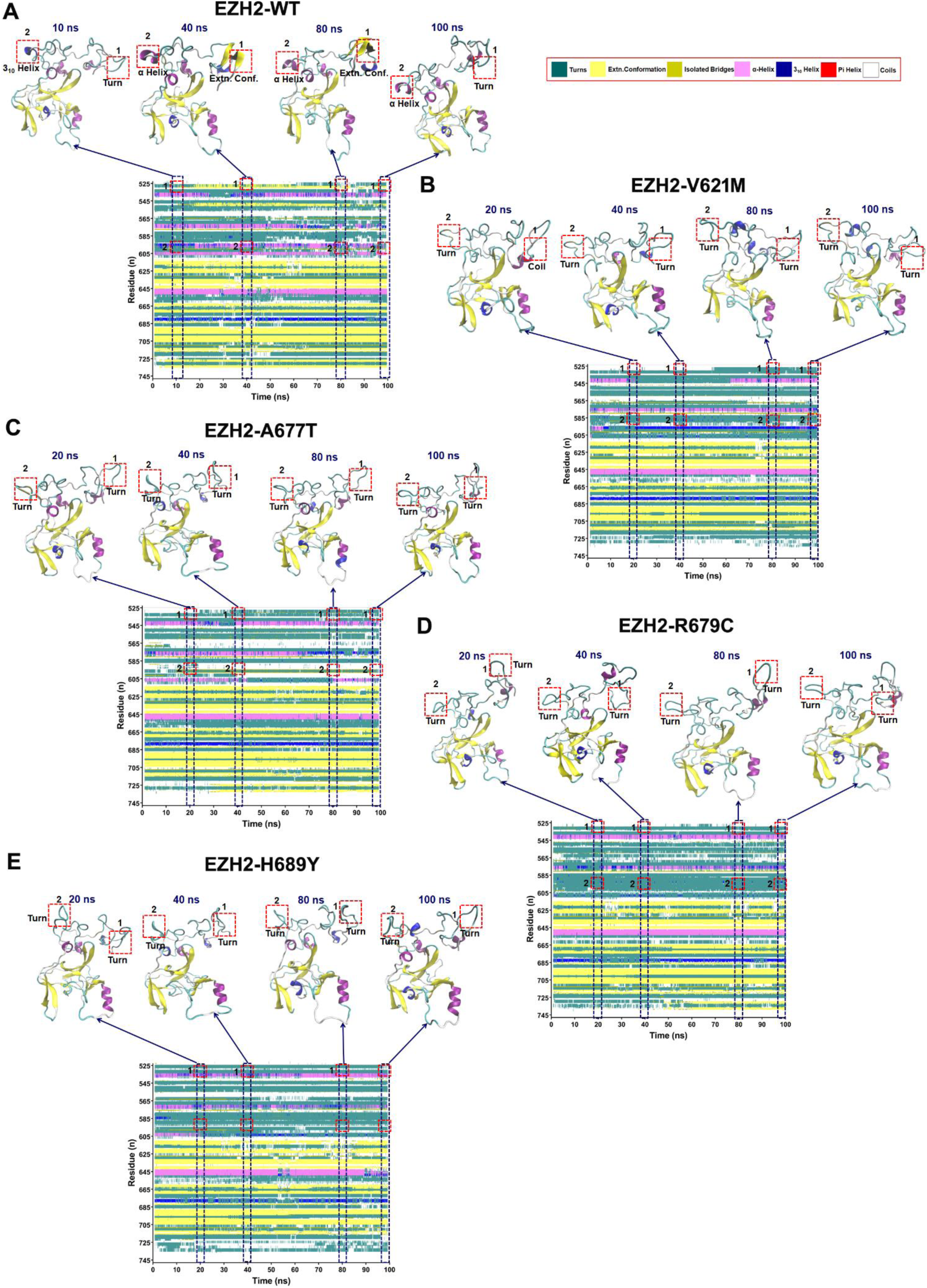
Secondary structure plots of EZH2-WT, EZH2-V621M, EZH2-A677T, EZH2-R679C and EZH2-H689Y during the 100 ns molecular dynamics simulation time period (**A, B, C, D and E**, respectively) reported through VMD timeline plugin. The significant secondary structure changes in the regions of 525-535 and 595-605 were depicted and highlighted at 20, 40, 80 and 100 ns with red dotted lines.

The structure of V621M has shown the partial structural change of turns to coil in the region of 526-530 amino acids up to 18ns further, the extended conformers has been replaced to turns in the same region throughout the 50ns of MDS. There is another conformational drift of 310 helixes and α helixes to turns in the region of 595-600 amino acids throughout the 50ns of MDS in comparison with wild-type EZH2 (Fig. 4B).

The mutant structure A677T has shown a conformational drift of extended conformer to turns with minor fluctuations to coils in the region of 526-530 amino acids throughout the 50ns of MDS. Another change of α helixes to turns with minor fluctuations has been found in the region of 595-600 amino acids throughout the 50ns of MDS. These changes were represented at 10ns, 20ns, 40ns and 50ns time-scale points (Fig. 4C). In addition, the MTs of R679C and H689Y have shown a conformational transition of extended conformers to turns and coils in the region of 526-530 amino acids up to 50ns of MDS. There is also another conformational drift of α helixes to turns in the region of 595-600 amino acids throughout the 50ns of MDS in comparison with wild-type structure of EZH2 (Fig. 4D, E).

The fluctuations of number of amino acids involved in the formation secondary structure elements were quantified for wild type and mutant structures during the 50ns of time scale period and were depicted in Figure 5A. The number of residues forming each secondary structure unit in the wild type and mutant structures was computed at every 10ns and the mean±SD values were plotted in bar graph (Fig. 5B). These statistical analyses demonstrate that mutants of R679C and H689Y have shown more significant increase of number of residues forming turns with p<0.001 while mutant of V621M has shown increase of turns with p<0.05 when compared to reported values in wild type structure. β sheets were found to be more significantly decreased in MTs of V621M and H689Y (p<0.001) and MTs of A677T and R679C were also shown significant increase with p<0.05 and p<0.01, respectively.

**Figure 5.**
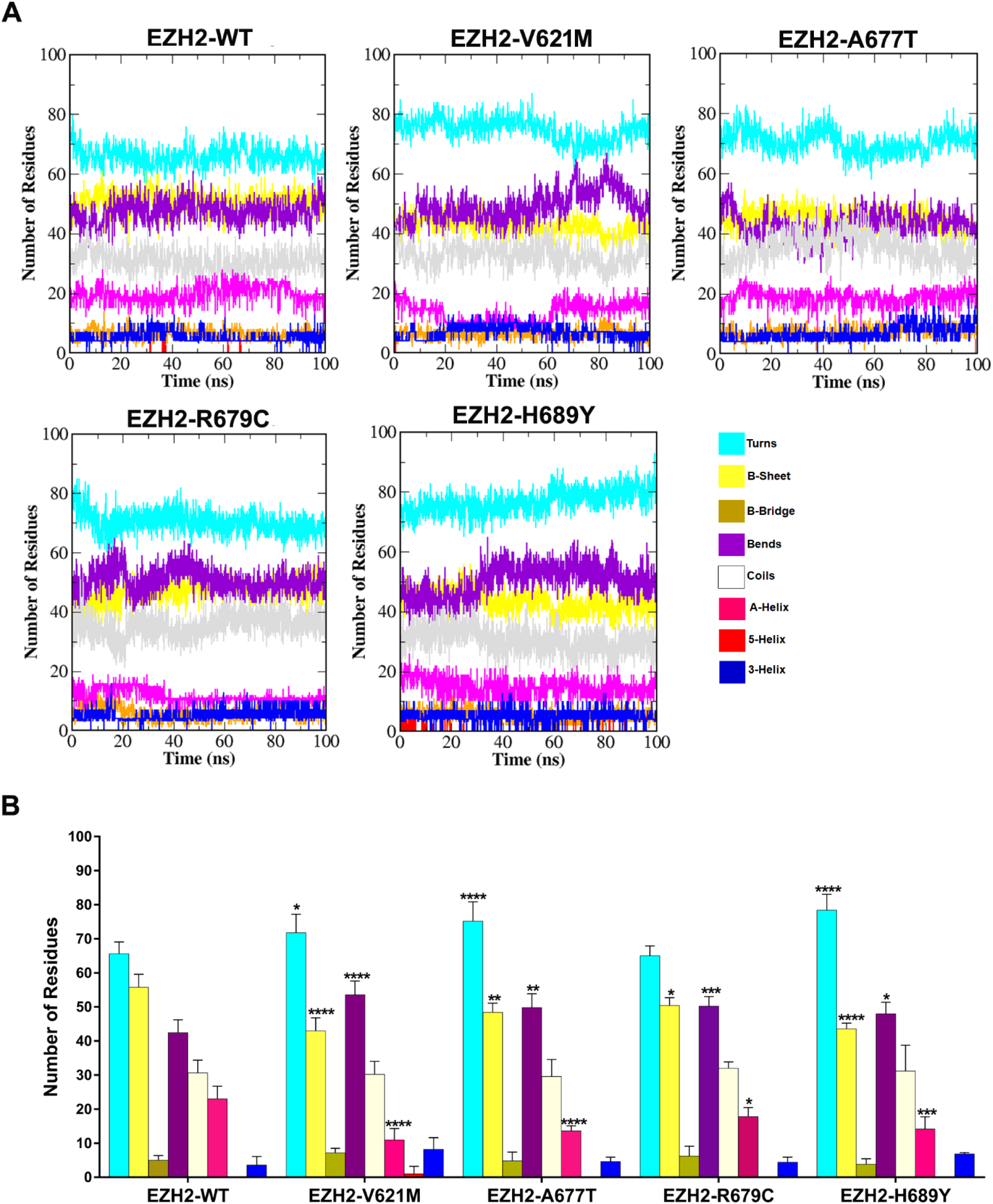
Secondary structure fluctuations of EZH2-WT, EZH2-V621M, EZH2-A677T, EZH2-R679C and EZH2-H689Y during 100 ns of time period of MD simulations through DSSP program **(A**). A number of amino acids involved in the formation various secondary structure units in EZH2-WT, EZH2-V621M, EZH2-A677T, EZH2-R679C and EZH2-H689Y. Error bars indicate SD from three replicates. The groups that have assigned **** (p<0.001), *** (p<0.005), **(p<0.01) and * (p<0.05) were statistically significant when compared to control.

The MTs of V621M and A677T have shown more significant increase of bends with p<0.001 and p<0.005, respectively and the MTs of R679C and H689Y were also shown significant increase with p<0.01 and p<0.05, respectively. In the same way, α helixes were more significantly decreased in MTs, V621M, R679C (p>0.001), A677T (p>0.05) and H689Y (p>0.005) when compared to reported values in WT structure. Hence it is obvious that the mutants of EZH2 have shown significant level of secondary structure alterations compared to wild type EZH2 (Fig. 5B).

The inter residue-residue contacts and fluctuations of pair-wise amino acid distance in WT and MT structures were characterized through distance map analysis. The results enumerate the variations of residue interactions among the MT structure in comparison with WT structure throughout the simulation time period. The distance plot analysis demonstrates that the three considerable pinpoint pair-wise locations in the WT structure, that has been found to be significant variations in the pair-wise distances among the MTs. The first distance pattern location in the WT structure the residues 15-25 as a turn and 310 helixes conformation with residues of 205-215 anti-parallel sheet in WT structure has significantly varied in MT structures, H689Y, A677T and V621M and the MT structure, R679C has not shown considerable variation.

The second distance pattern residues, 50-62 as a α helix conformation with 220-226 of N-terminal turns and the third distance pattern residues, in the WT structure and the third distance pattern residues 220-226 with 190-200. All these changes were represented as rectangular boxes in Figure S4. Hence, the overall distance pattern observed in the WT structure is a vital conformation for functional catalysis of protein. The MT structures have been found to be undergone conformational rearrangement and leads to functional impairment. Thus, the overall results have shown good agreements with Cα atom fluctuations depicted in RMSF plot.

The final refined model of WT and MT structures after 100ns of MD simulations were subjected for Ramachandran plot calculations through PROCHECK server to verify the reliability the protein structure. The results showed that the MTs of H689Y, R679C, A677T, and V621M have the residues plotted in the most favorable region of Ramachandran plot, 75.7, 75.7, 76.70 and 79.70%, respectively when compared to 80.7% for WT structure. Therefore, the total number of residues plotted in each region (most favorable, additionally allowed, generously allowed and disallowed regions) of Ramachandran plot for WT and MT structures were depicted in a Figure S5B and summarized as a list in a Figure S5B

### 3.5. Molecular dynamics of Protein-peptide complexes

Protein-peptide docking studies were employed to analyze the interactions of EZH2-WT and its mutants with H3K27me0 peptide and to show the effect of corresponding mutations in the binding affinity and H-bond interactions with H3K27me0. The docking program has generated 10 different docked poses in which the best fitted conformation was selected for each protein. The results of docking analysis demonstrate that the MTs of A677T and H689Y have shown lower binding affinity with -6.3 and -6.1 kcal/mol, respectively when compared to -7.8 kcal/mol for EZH2-WT structure (Fig. 6A) and molecular interactions of H3K27me0 peptide with WT and MT structures were also depicted (Fig. 6B-F).

**Figure 6.**
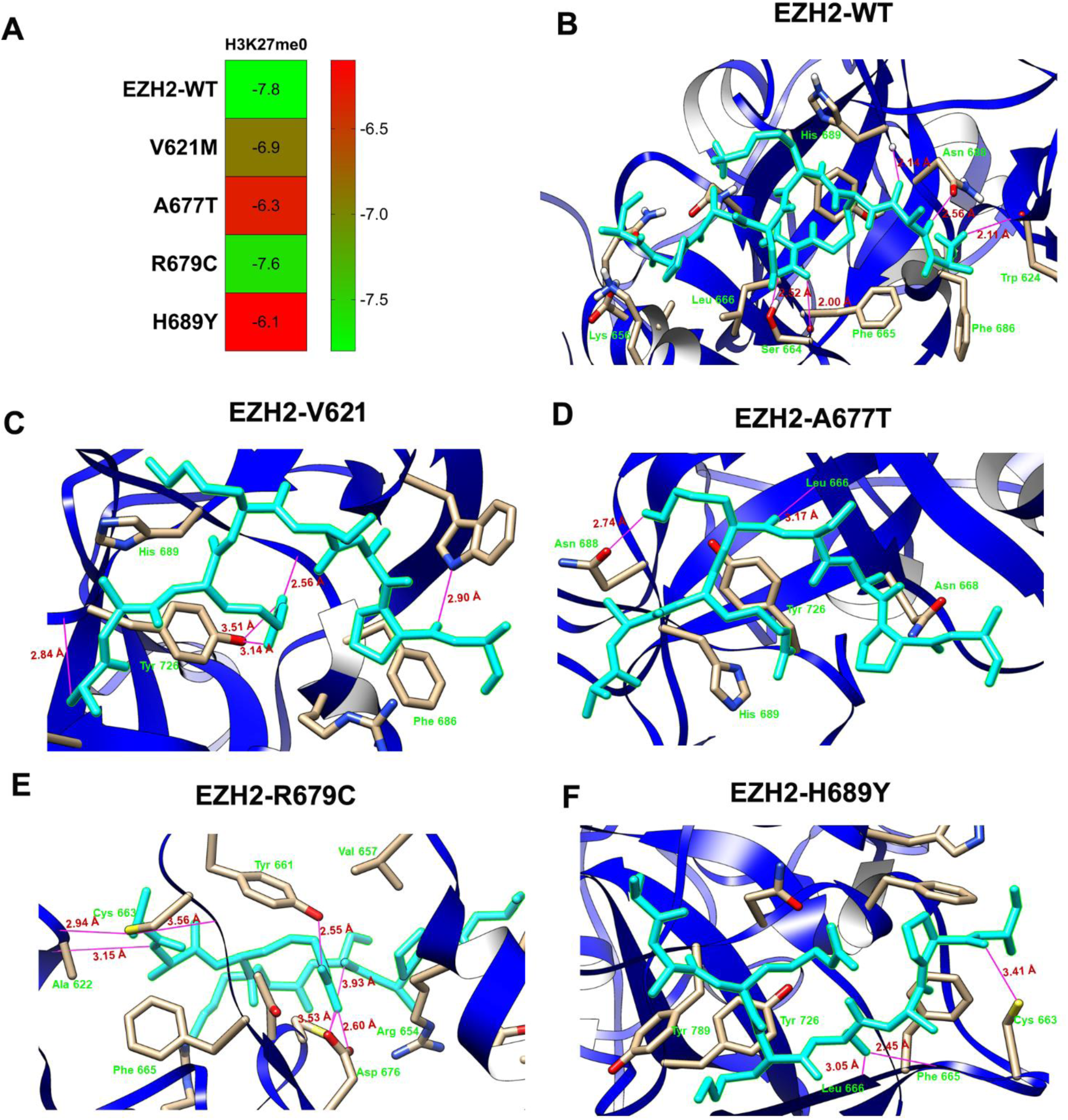
Molecular interactions of H3K27me0 with active site residues of EZH2-WT, EZH2-V621M, EZH2-A677T, EZH2-R679C and EZH2-H689Y (**A, B, C, D and E**, respectively) after 50 ns of MD simulations. Heat map representation of binding affinity of H3K27me0 with binding pocket of WT and MT structures.

Binding orientation of H3K27me0 peptide with EZH2-WT showed the formation of five H-bonds in which three H bonds were formed with the residues Trp 624, Asn 688 and His 689, respectively while the other two H bonds were with the residue Ser 667. (Fig. 6B). The MT structures, A677T and H689Y have shown significant changes in H-bonding in comparison with WT structure. In the A677T, the H3K27me0 has shown only two H-bonds with Leu666 and Asn 688, respectively (Fig. 6D). While in the H689Y, three H-bonds were formed with Cys663, Phe665 and Leu666, respectively (Fig. 6F). However, the other two MTs V621M and R679C have not shown obvious changes in H-bond number when compared to WT structure (Fig. 6C, E).

Further, Protein-peptide docked complexes such as EZH2-WT-H3K27me0, V621M-H3K27me0, A677T-H3K27me0, R679-H3K27me0 and H689Y-H3K27me0 were subjected to the 50ns of MD simulations for comprehensive studies on the stability and dynamics of the complex. The MDS were carried out to investigate the stability mode of interactions of protein-peptide complexes by means of RMSD, RMSF, H-bond profile and secondary structure.

The results of calculated RMSD of protein and protein-peptide complexes with respect to the initial model and during 100ns of MDS were depicted in the form of heatmap (Fig. S6). The results demonstrate that the mutant structures, A677T, R679C and H689Y binds with H3K27me0 peptide has shown higher level of fluctuations of RMSD with maximum value of 2.0, 1.65 and 1.67 nm, respectively when compared to peptide free MTs. However, the V621M mutant and wild-type EZH2 in complex with H3K27me0 peptide have not shown significant variations in RMSD with 0.798nm and 0.579nm when compared to peptide-free V621M mutant and the wild type, respectively. We also plotted the fluctuations of RMSD values for mutant and wild type complexes with H3K27me0 peptide. From this plot it is evidenced that MT complex A677T-H3K27me0 has shown the minor fluctuations of rmsd within the range of 0-0.25 nm in the non-diagonal region followed by sharp increase up to 1.95 nm after 39 ns and finally maintains equilibrium to the 50 ns of MD simulations. In the same manner MT complexes R679-H3K27me0 and H689Y-H3K27me0 have shown minor fluctuations of rmsd with average 0.60 and 0.30 nm in the non-diagonal region, respectively. However, the MT complex V621M-H3K27me0 has shown average rmsd of 0.35 nm throughout the 50 ns of MD simulations similar with the levels in WT-H3K27me0 complex (Fig. S7A).

The results of H-bonding profile demonstrate that the WT structure has shown maximum of seven H-bond interactions with H3K27me0, among which four bonds were stable whereas three bonds were relatively weaker. The bonding profile and intensity throughout the 50ns of time period of MDS was reported in Figure 8A. The intensity of the H bond in the plot was quantified based on the continuity of the band throughout the 50 ns of MDS. Whereas MT structures of A677T, R679C and H689Y have shown significant reduction of H-bonds with H3K27me0 when compared to WT structure (Fig. B-E).

**Figure 7.**
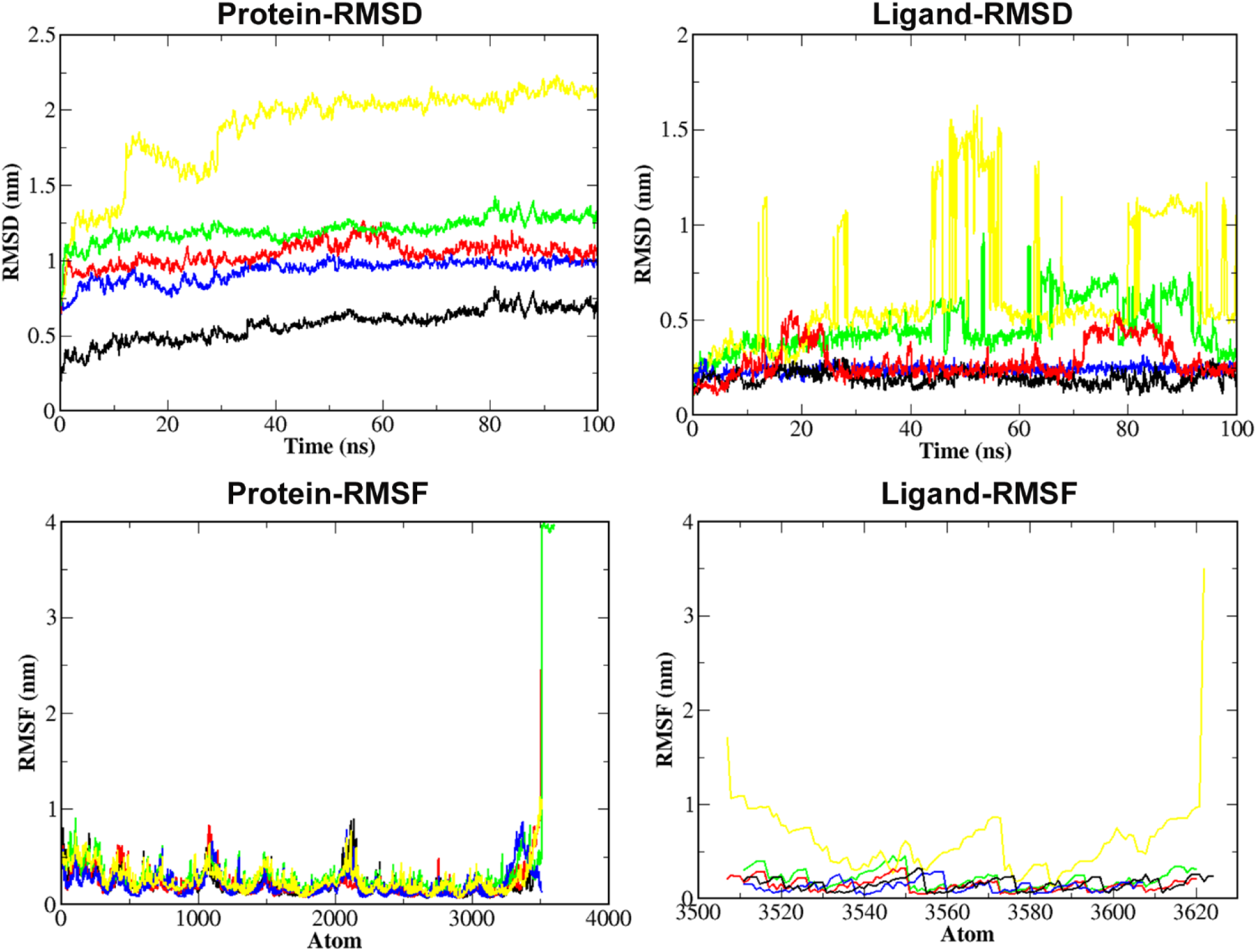
The backbone Cα atoms Root mean square fluctuations of EZH2-WT, EZH2-V621M, EZH2-A677T, EZH2-R679C and EZH2-H689Y during interaction with H3K27me0 peptide and the RMSF of H3K27me0 in the binding pocket of WT and MT structures.

**Figure 8.**
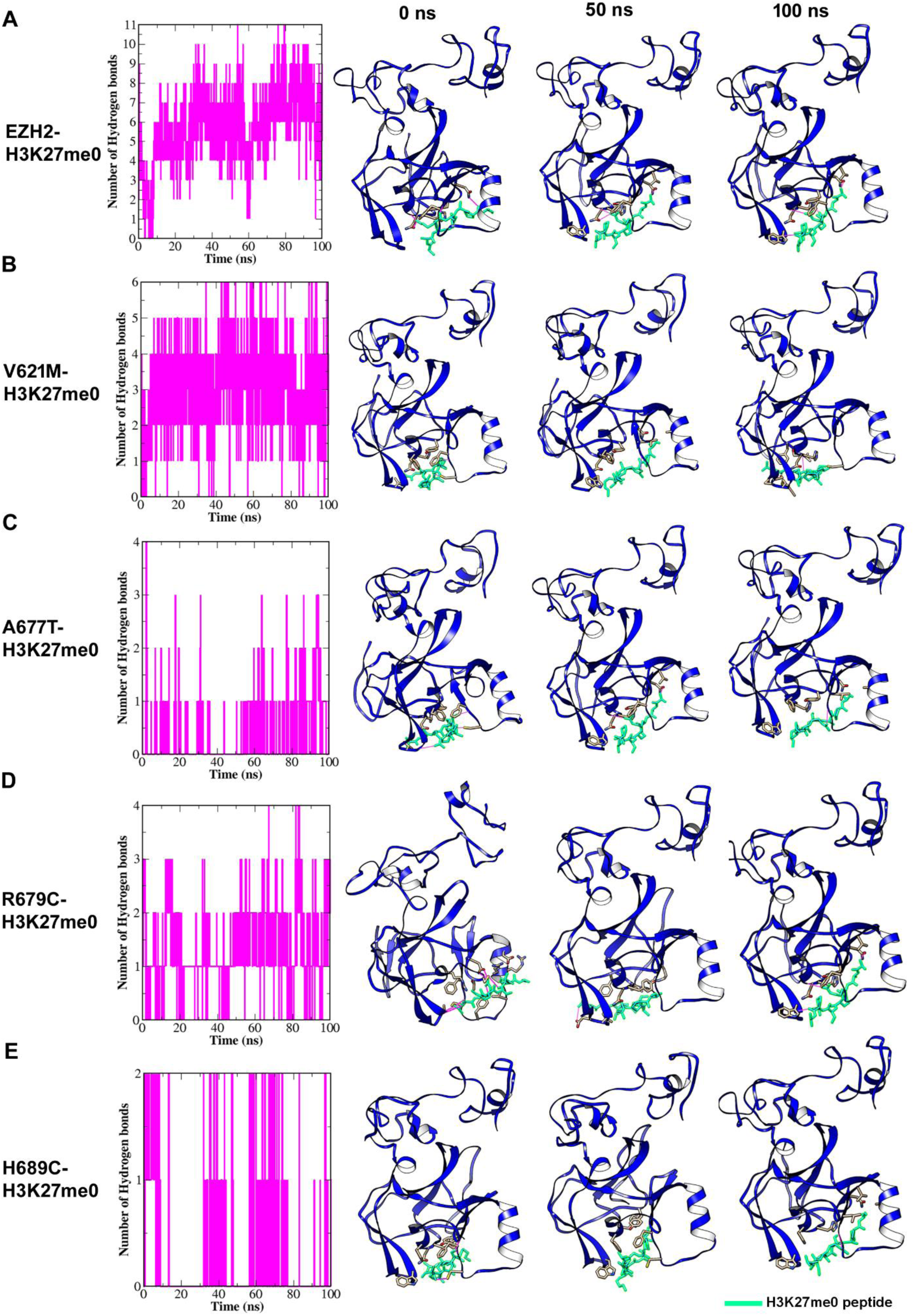
Hydrogen bond intensity between H3K27me0 and binding pocket residues of EZH2-WT, EZH2-V621M, EZH2-A677T, EZH2-R679C and EZH2-H689Y (**A, B, C, D and E**, respectively) during 100ns of time period of MDS with snapshots of binding poses at 0, 50 and 100 ns time period.

In WT-H3K27me0 complex, the peptide has shown constant and stable interactions with three H-bonds. The binding orientation and poses were captured at 0, 25 and 50 ns time period of MDS. The results showed that the peptide has shown stable conformation in the time period (Fig. 8A). In the MT-H3K27me0 complex the peptide has formed two stable H bonds with V621M and H689Y, and one stable H bond with A677T. In addition, the binding orientation and the peptide conformation have significantly changed when compared to WT-H3K27me0 (Fig. 8B, C, E). In the case of R679C-H3K27me0 complex, though the scale of H bond plot has shown the maximum of 10 H bonds, there were only three H bonds that were stable and showed strong intensity throughout the 50 ns of MD simulations. Hence, we conclude that the H3K27me0 peptide has shown fewer binding affinities and unstable binding conformations with the EZH2 mutants.

### 3.6. Principal Component analysis (PCA)

In order to characterize the large-scale motions of MT and WT proteins, we performed a principal component analysis (PCA) of covariance matrix from the MD trajectory. The original space of correlated variables has been transformed to a reduced space of independent variables (viz., principal components or eigen vectors) through principal component analysis. The PCA identifies the amplitude and direction of dominant protein motions by projecting the trajectories in a reduced subspace.

From the cluster of 678 eigen vectors corresponding for Cα atoms of the WT & MT proteins, in this first twenty eigen vectors were selected and plotted against the corresponding eigen values (Fig. S9). The plot demonstrates that the first two eigen vectors showed larger eigen values for both WT and MT proteins. Hence, the projection of trajectories of WT and MT proteins in the phase space along the first two principal components, PC1 and PC2 were plotted (Fig. 9). From these plots it is very obvious that the model of WT was well defined than in the models of MTs. The MT structures occupied larger region of space along the first principal component (PC1) when compared with WT structure. The Cα atom fluctuations of principal components were analyzed to verify the atomic motions of WT & MT structures. The PC analysis of WT structure has shown the most pronounced atomic motions of coils in the amino acid region of 25-30 and the α Helices in the amino acid region of 125-140 in the PC1 and PC2, respectively (Fig. 9A). The MT structures V621M, A677T and R679C have shown higher atomic motions of N-terminal coils in PC1 (Fig. 9B, C, D). Whereas the MT structure H689Y has shown higher atomic motions of C-terminal coils in the PC1 (Fig. 9E).

**Figure 9.**
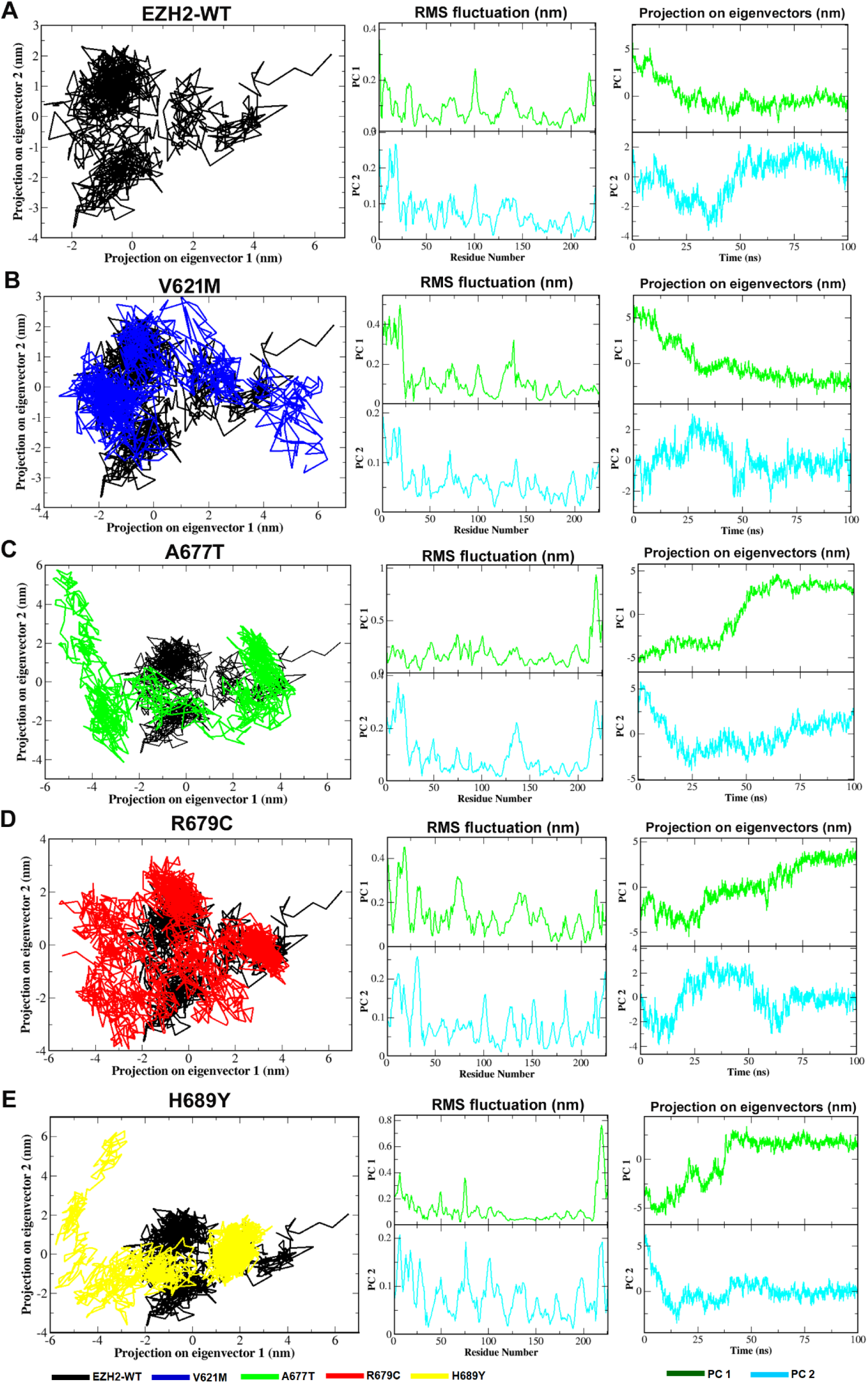
Projection of motion of the protein in the phase and RMS fluctuations of first two principal components and structure alignments of EZH2-WT(A) and mutant structures, V621M (B), A677T (C), R677T (D) and H689Y (E).

Further, the overall flexibility of WT, MTs and complexes with H3K27me0 peptide was investigated by the trace of diagonalized covariance matrix of Cα atoms. The resulted covariance plots have portrayed the positive and negative limits of values, which related to the motion of atoms occurring along the same direction and the opposite direction, respectively. The highly anti-correlated motions were supporting the stability of the model and found to be WT structure when compared to the MT structure. The trace values for WT, V621M, A677T, R679C and H689Y were found to be 6.51, 7.73, 13.54, 11.58 and 14.25 nm2, respectively (Fig. S10A).

Therefore, the covariance matrix for the complex of mutant and H3K27me0 has demonstrated that the binding of H3K27me0 with mutant has induced higher level of structural flexibility when compared to wild-type protein and native peptide free MTs. The resulted trace values of covariance matrix for EZH2-WT-H3K27me0, V621M-H3K27me0, A677T-H3K27me0, R679-H3K27me0 and H689Y-H3K27me0 were found to be 4.89, 12.80, 160.72, 29.87 and 39.57 nm2, respectively (Fig. S10 B). Hence, the higher trace value for mutants and its complexes indicated increased structural flexibility when compared to wild-type.

### 3.7. Residue interaction network analysis

RIN analysis was performed to explore the characterization of molecular interactions of EZH2 residues within the peptide binding pocket. The results of RIN plot analysis revealed a noticeable difference between WT and MT structures at various nano second time periods.

The WT structure at 0 ns has shown the compact of different bonding network such as H-bond, Van der Waals, and PiPi-stack interactions. There are four VDW interactions at the residues 686-624, 676-666, 726-666 and 726-689, two Pi-Pi stack interactions at the residues 686-626 and 726-689, and one H-bond interaction between the residues 726-689. At 25 and 50 ns, WT structure has shown the bonding network of three VDW between the residues 676-666, 679-661 and 726-688, respectively, and one H bond and PiPi stack between the residues 726-689. One H bond between the residues 688-624 has confined to only 25 ns of time period. (Fig. 10A). The interaction network between these residues formed the unique configuration that is allowed to interact with H3K27me0 peptide.

**Figure 10.**
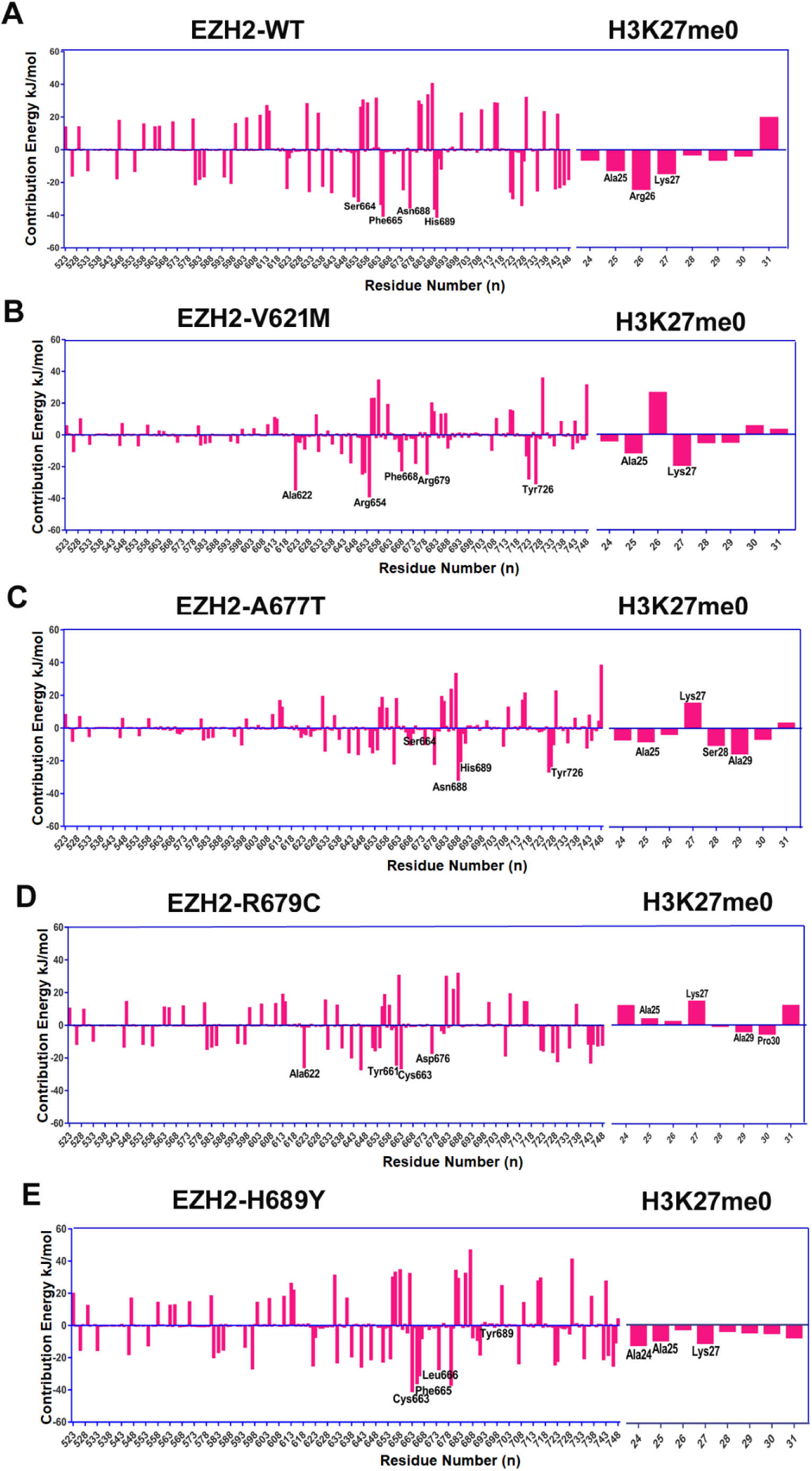
Comparison of residue interaction network (RINs) of binding pocket residues of WT and MT structures, V621M, A677T, R679C and H689Y at 0 ns, 50 ns and 100 ns time period of MD simulations.

The RIN analysis has shown that the mutant residues changed the interaction network when compared to the wild type. The MT structure V621M has shown the interaction profile of two VDW and PiPi-stack interactions between the residues 624-686 and 726-689, at 0 ns of time period, respectively. At 25 and 50 ns, the V621M has shown one H bond and VDW interaction between the residues 726-689 and one Pi-Pi stack confined to only 25 ns of time period (Fig. 10B). The MT structure A677T has shown four VDW interactions at the residues 676-666, 679-661, 688-726 and 689-726, two H bonds between the residues 689-726 and 688-624, and one PiPistack interaction between the residues 689-726 at 0 ns of MD simulation. At 25 ns, A677T has shown the absence of one H bond and VDW interaction between the residues 624-688 and 661-679, respectively, but one new VDW interaction has formed between the residue 622-688 when compared to interactions at 0 ns. At 50 ns, A677T loses of two H bonds between the residues, 624-688 and 689-726, respectively, and one VDW interaction between the residue 688-726 when compared to the interactions at 0 ns (Fig. 10C).

A bonding network of four VDW and one PiPistack interactions in the MT structure, R679C and one H bond, VDW and PiPi stack interactions in the MT structure H689Y were observed at 0 ns. The significant reduction of bonds was observed at 25 and 50 ns time period of MD simulations (Fig. 10D, E) Hence the RIN analysis reveals that the dynamic orientation and bonding network has significantly changed in the MT structures.

## 4. Discussion

Screening and characterization of non-synonymous SNPs in the coding region of ezh2 gene involved in Weaver Syndrome was the fundamental objective of the current research. We focused on four nsSNPs including V621M, A677T, R679C and H689 with deleterious effects on EZH2. Earlier biochemical studies have been reported that the aforementioned nsSNPs affected enzymatic activity of EZH2 and lower the methylation level of H3K27. The de nova missense mutation V621M was identified and found to be expressed in growth plate chondrocytes. The mutant protein has shown the decreased activity of histone H3K27 methylation in the in vitro assay and in the mouse, model using CRISPR/Cas9 [39]. The mutants of A677T, R679C and H689Y were found to be associated with neuroblastoma, acute lymphoblastic leukemia and lymphoma in the WS patients [40] and shows significant reduction of histone H3K27 methylation when compared to wild type protein [18].

Keeping in mind of the physiological relevance of these mutants, the computational methods, docking and molecular dynamics simulations were employed to elucidate the structural variations and effects of the mutative amino acids in the interactions with their common substrate H3K27me0. The MD trajectories of RMSD, RMSF, Rg, SASA and density plot analysis demonstrate that the mutants of H689Y, R679C and A677T have shown significant variations when compared to WT structure. The protein-peptide docking reveals that the MTs of H689Y and A677T have shown less binding affinity to H3K27me0 with -6.1 and -6.3 kcal/mol, respectively. The protein docking was supported by RMSD and RMSF analysis of complexes. The MTs of A677T and H689Y have shown higher RMSD values with 2.0nm and 1.67nm, respectively, when interacting with H3K27me0 peptide. The H3K27me0 peptide has shown higher RMSF values in the binding pocket of MT structures of A677T and H689Y during their interactions, which led to weak binding affinity. It is also evidenced by H-bond profile when compared to wild type (Fig.7, 8). The peptide binding pocket in the SET-domain of EZH2 has composed of 310 helix, ext. conformations and turns around the region of 675-700 amino acids. The secondary structure plot of MT structures during treated with H3K27me0 peptide has shown that the denature of 310 helixes around the region of 680-690 amino acids in the MT structures of A677T and H689Y when compared to wild type (Fig. S9). The amino acids Arg 685, Phe 686, Ala 687, Asn 688 and His 689 stabilize the formation of 310 helix. In the wild type structure, the amino acids Phe 686, Asn 688 and His 689 have formed stable H-bonds with the H3K27me0 peptide and anchored for the transfer of methyl group from cofactor S-adenosyl methionine (SAM). In the MT structures of A677T and H689Y, the H3K27me0 has formed no H-bond interactions with the aforementioned amino acids due to the destabilization of 310 helix when compared to wild type.

Principal component analysis (PCA) is a multivariate statistical technique for protein dynamics and can be used to study the molecular motions in the smallest spatial scale [41, 42]. PC analysis of EZH-WT structure has shown more profound motions of coils in the PC1 and α Helixes in PC2. Whereas the PC1 of MT structures V621M, A677T and R679C have shown higher atomic motions of N-terminal coil and H689Y has shown profound motions of C-terminal coil as exemplified by RMS fluctuations PC1 and PC2.The covariance analysis of MTs and MT-H3K27me0 complexes has shown correlated motions that destabilize their own structures, when compared to WT structure. The MT structures of A677T and H689Y have shown higher correlation motions with the trace values of 160.72 and 39.57 nm2, during their interactions with H3K27me0 that destabilize the complexes with lower binding affinities. Overall, the domain network and dynamic amino acid orientations in the binding pocket strongly influence the peptide recognition [43]. Hence, we observed the considerable correlations between the computational and in vitro experiments. As exemplified by our data, EZH2 was an emerging epigenetic repressive target and its SET-domain was a protein-protein interactive domain. Our work lay a frontier for the discovery of small molecule inhibitors using computer-aided drug design methods against Weaver syndrome.

## 5. Conclusions

In conclusion, the molecular modeling and dynamics methods offer an opportunity for comprehensive structural and functional characterization of EZH2 mutants of V621M, A677T, R679C, and H689Y. MD simulation of aforementioned mutants and their complexes with H3K27me0 were employed to decipher the stability and effects of corresponding amino acid replacements in H-bonding profiles. The RMSD, Rg, SASA and density plots revealed their deleterious nature of A677T, R679C and H689Y mutants. H-bond profile and covariance analysis revealed that mutants of A677T and H689Y have weaker H-bond interactions with H3K27me0 and higher covariance trace value due to the destabilization of α-helixes within the peptide binding region. Overall, the computational analysis provides a thorough insight into the destabilizing mechanisms of EZH2 nsSNPs involved in Weaver syndrome and is helpful for translational research of EZH2-mutant-based therapeutic strategies against genetic disorders.

## Author Contributions

Conceptualization, Tirumalasetty Muni Babu, Danian Qin and Chengyang Huang; Data curation, Vivek Keshri; Formal analysis, Tirumalasetty Muni Babu and Vivek Keshri; Funding acquisition, Danian Qin and Chengyang Huang; Investigation, Tirumalasetty Muni Babu, Vivek Keshri and Cuicui Yu; Methodology, Tirumalasetty Muni Babu and Chengyang Huang; Project administration, Danian Qin and Chengyang Huang; Resources, Cuicui Yu; Software, Tirumalasetty Muni Babu and Vivek Keshri; Supervision, Danian Qin and Chengyang Huang; Validation, Cuicui Yu; Visualization, Tirumalasetty Muni Babu and Vivek Keshri; Writing – original draft, Tirumalasetty Muni Babu, Vivek Keshri and Chengyang Huang; Writing – review & editing, Danian Qin and Chengyang Huang.

## Acknowledgments

We are thankful to SUMC for providing research facilities. We are also grateful to the members of the Huang lab for helpful comments and discussion.

