## Supplemental Figures and Tables for "Structural dynamics of non-synonymous SNPs of histone methyltransferase EZH2 involved in Weaver Syndrome"

**Figure S1**

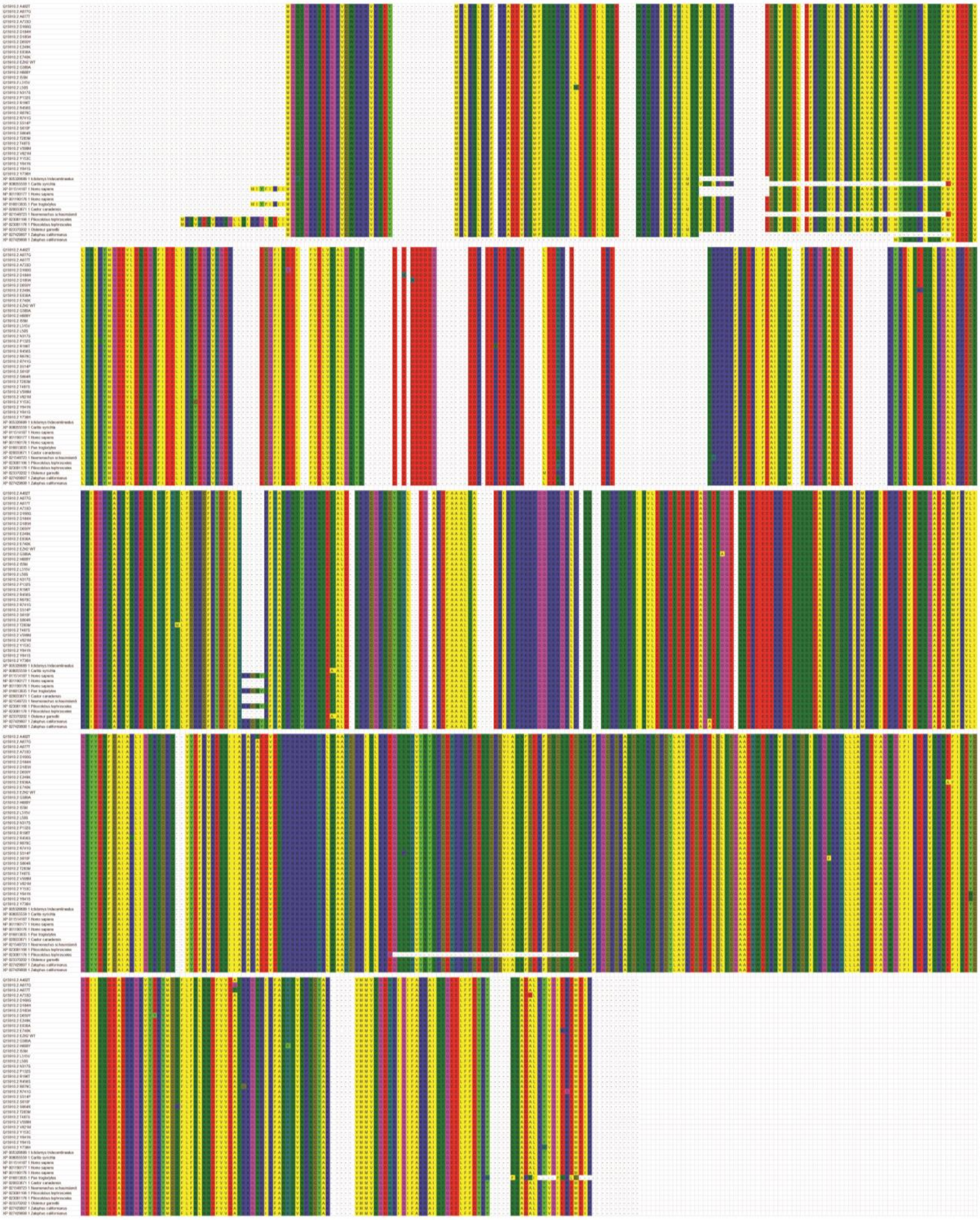

**Figure S2**

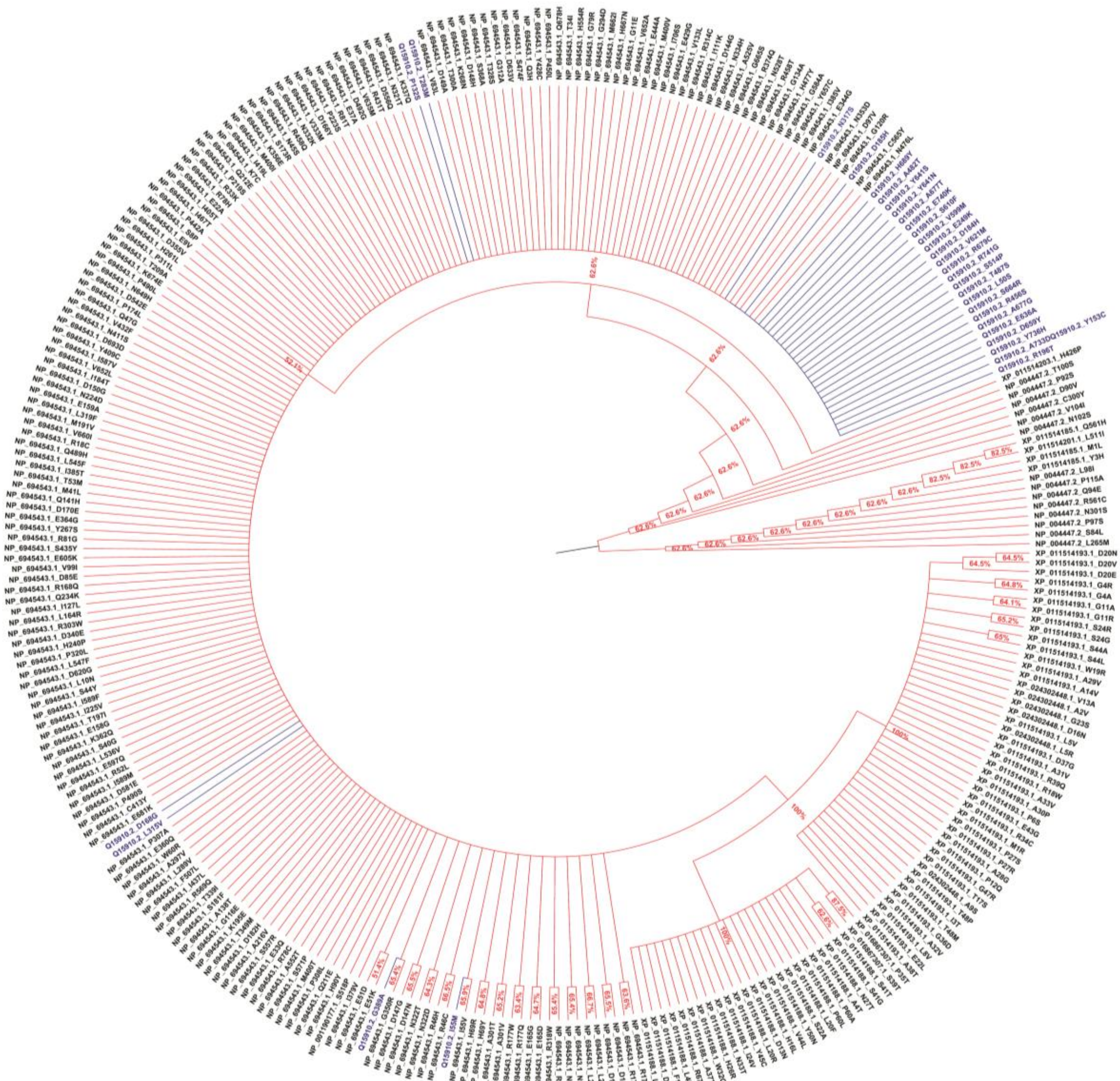

#### Figure S3

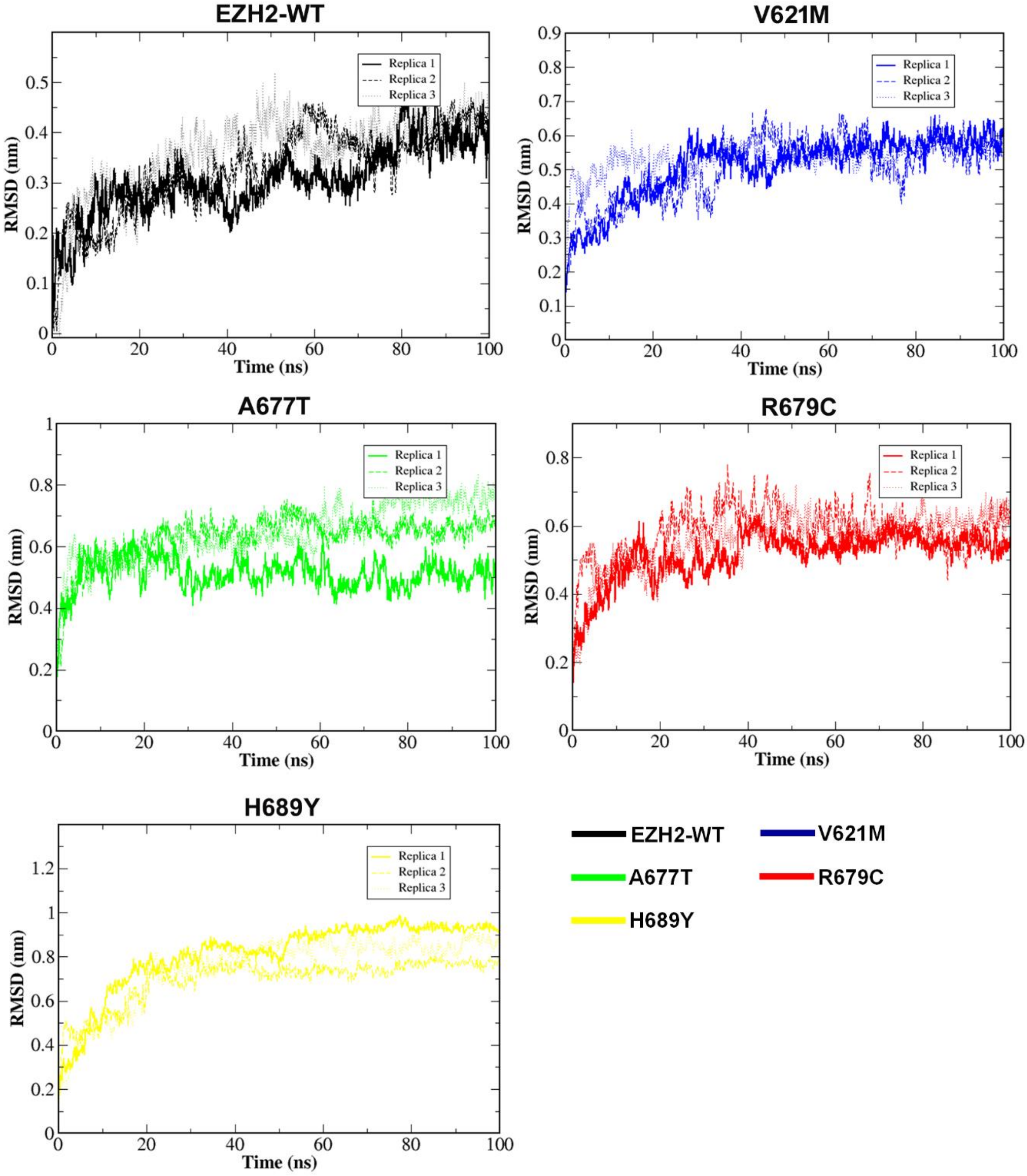
EZH2-WT
V621M
A677T
R679C
H689Y
RMSD (nm)
Time (ns)
Replica 1
Replica 2
Replica 3
EZH2-WT
V621M
A677T
R679C
H689Y

#### Figure S4

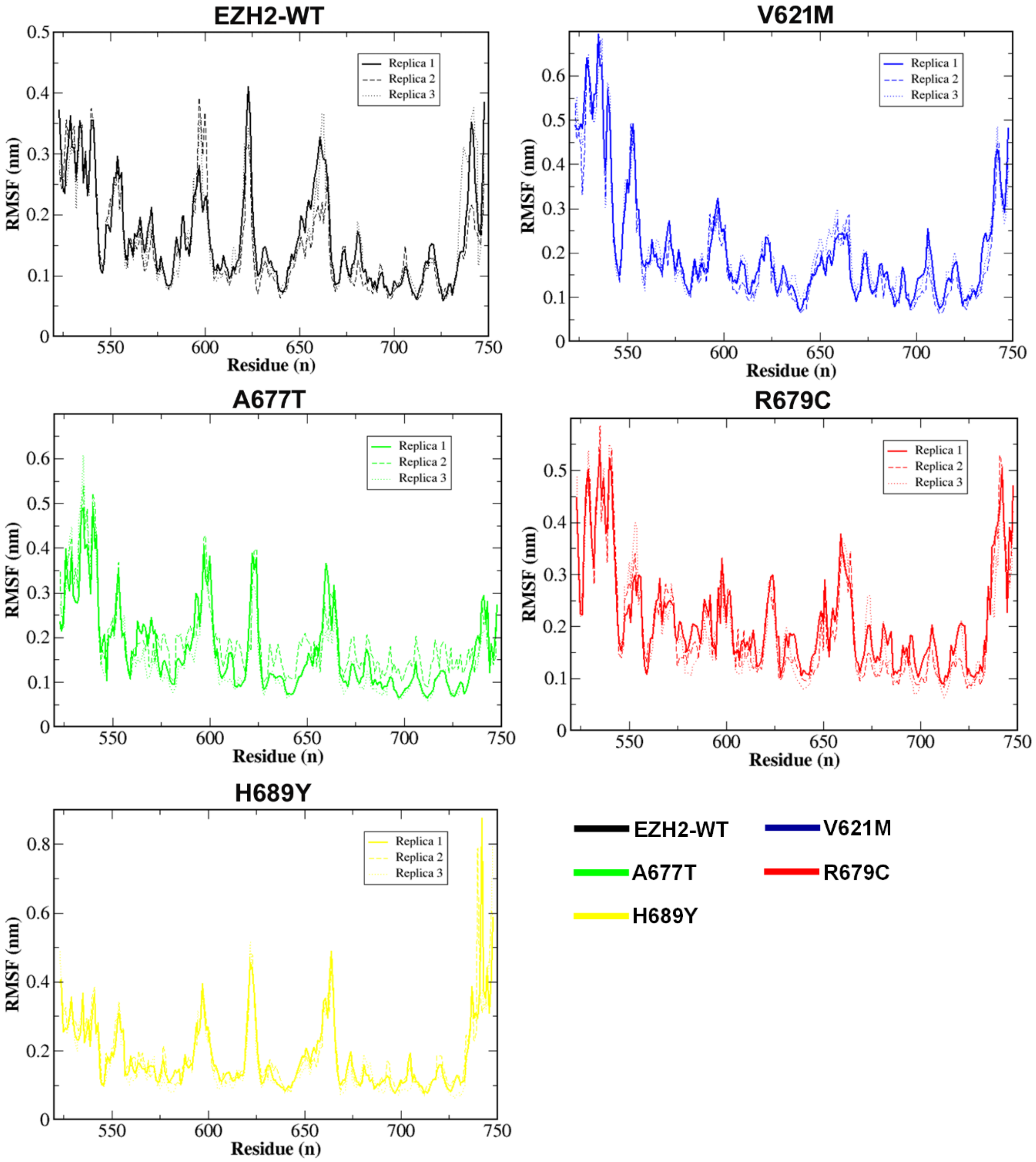
EZH2-WT
V621M
A677T
R679C
H689Y
RMSF (nm)
Residue (n)
Replica 1
Replica 2
Replica 3
EZH2-WT
V621M
A677T
R679C
H689Y

### Figure S5

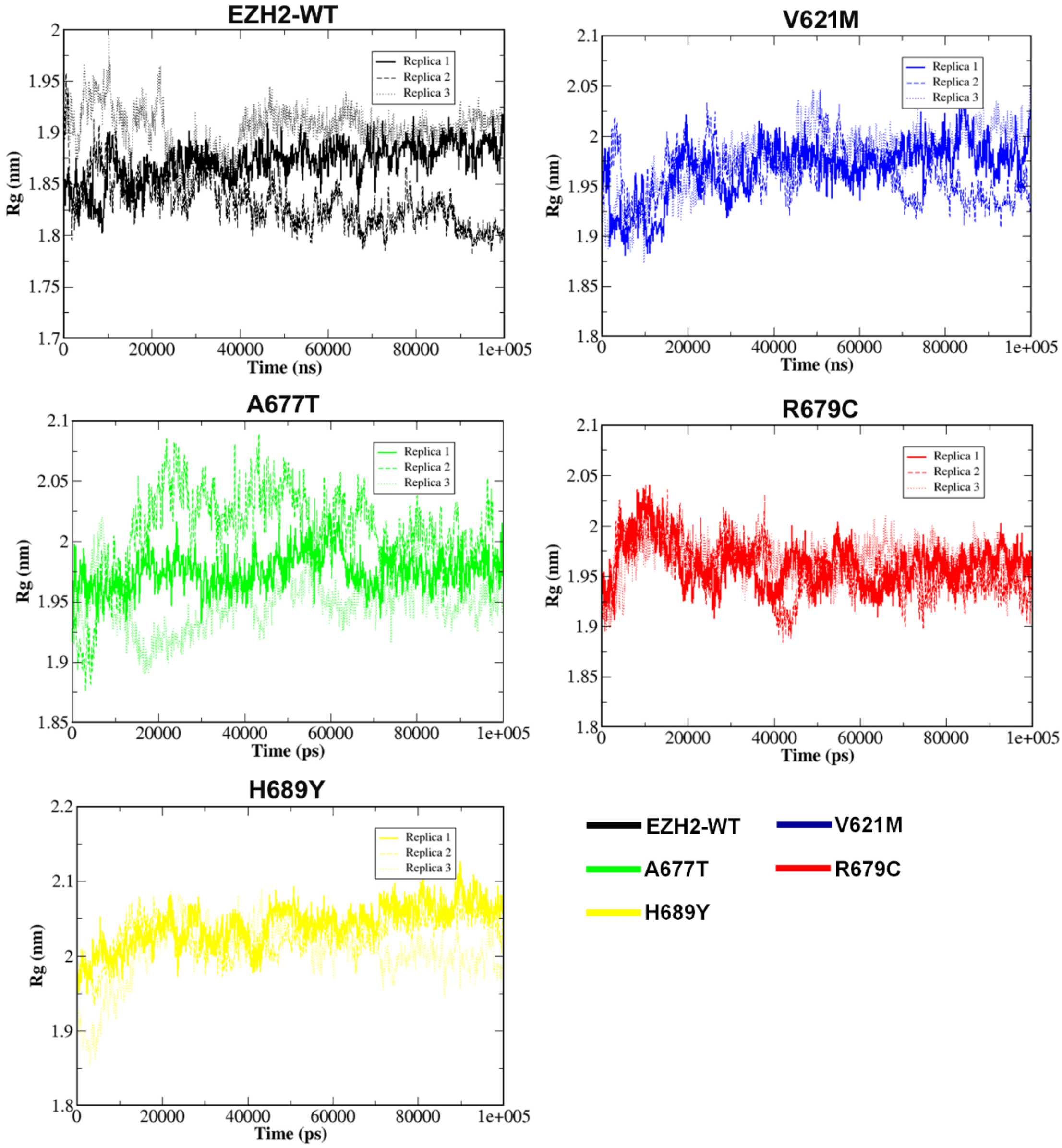
EZH2-WT
Replica 1
Replica 2
Replica 3
Rg (nm)
Time (ns)
V621M
Replica 1
Replica 2
Replica 3
Rg (nm)
Time (ns)
A677T
Replica 1
Replica 2
Replica 3
Rg (nm)
Time (ps)
R679C
Replica 1
Replica 2
Replica 3
Rg (nm)
Time (ps)
H689Y
Replica 1
Replica 2
Replica 3
Rg (nm)
Time (ps)
EZH2-WT
V621M
A677T
R679C
H689Y

#### Figure S6

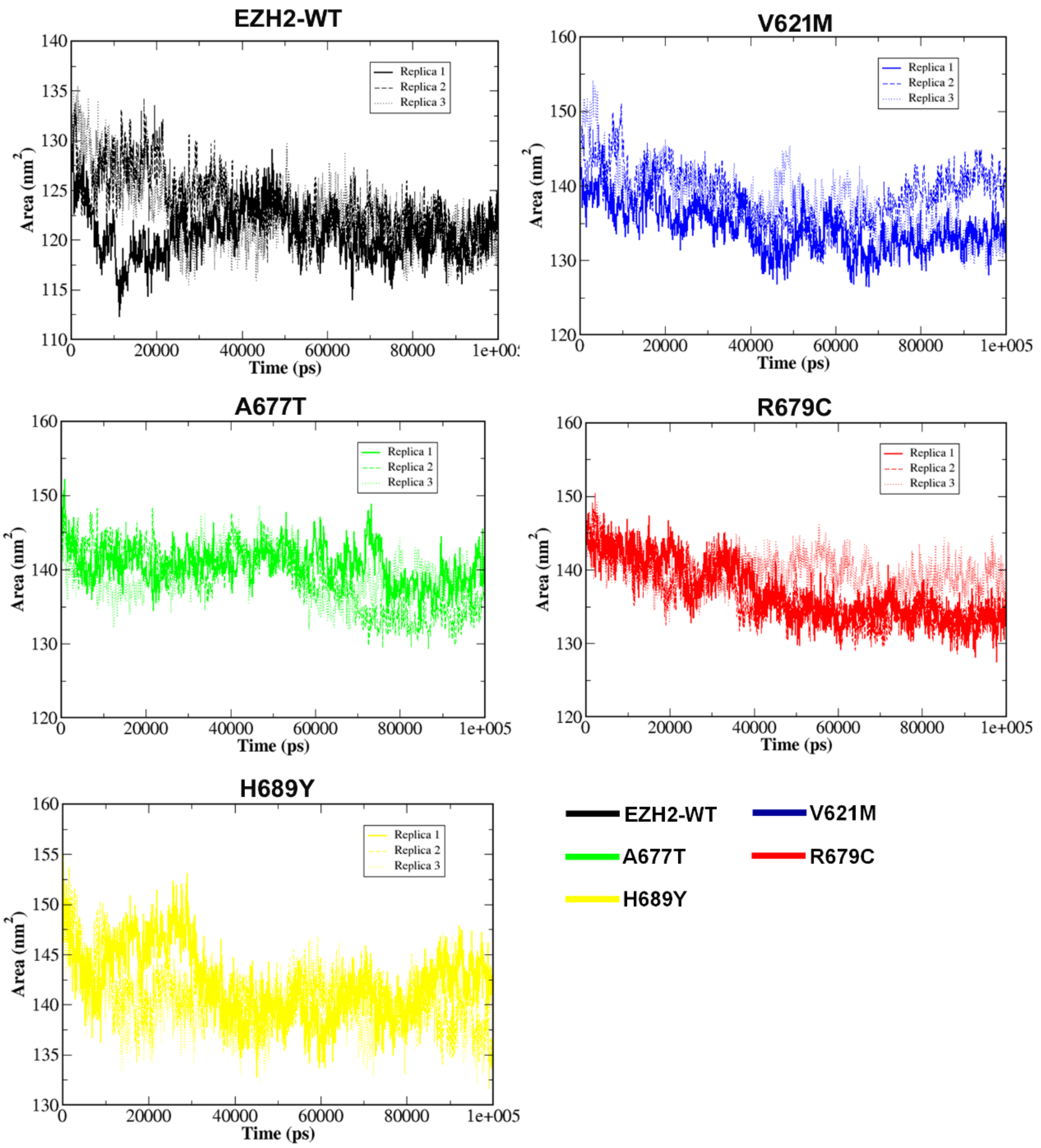

EZH2-WT
V621M
A677T
R679C
H689Y
Replica 1
Replica 2
Replica 3
Area (nm$^2$)
Time (ps)
0
20000
40000
60000
80000
1e+005
110
115
120
125
130
135
140
145
150
155
160
EZH2-WT
V621M
A677T
R679C
H689Y

### Figure S7

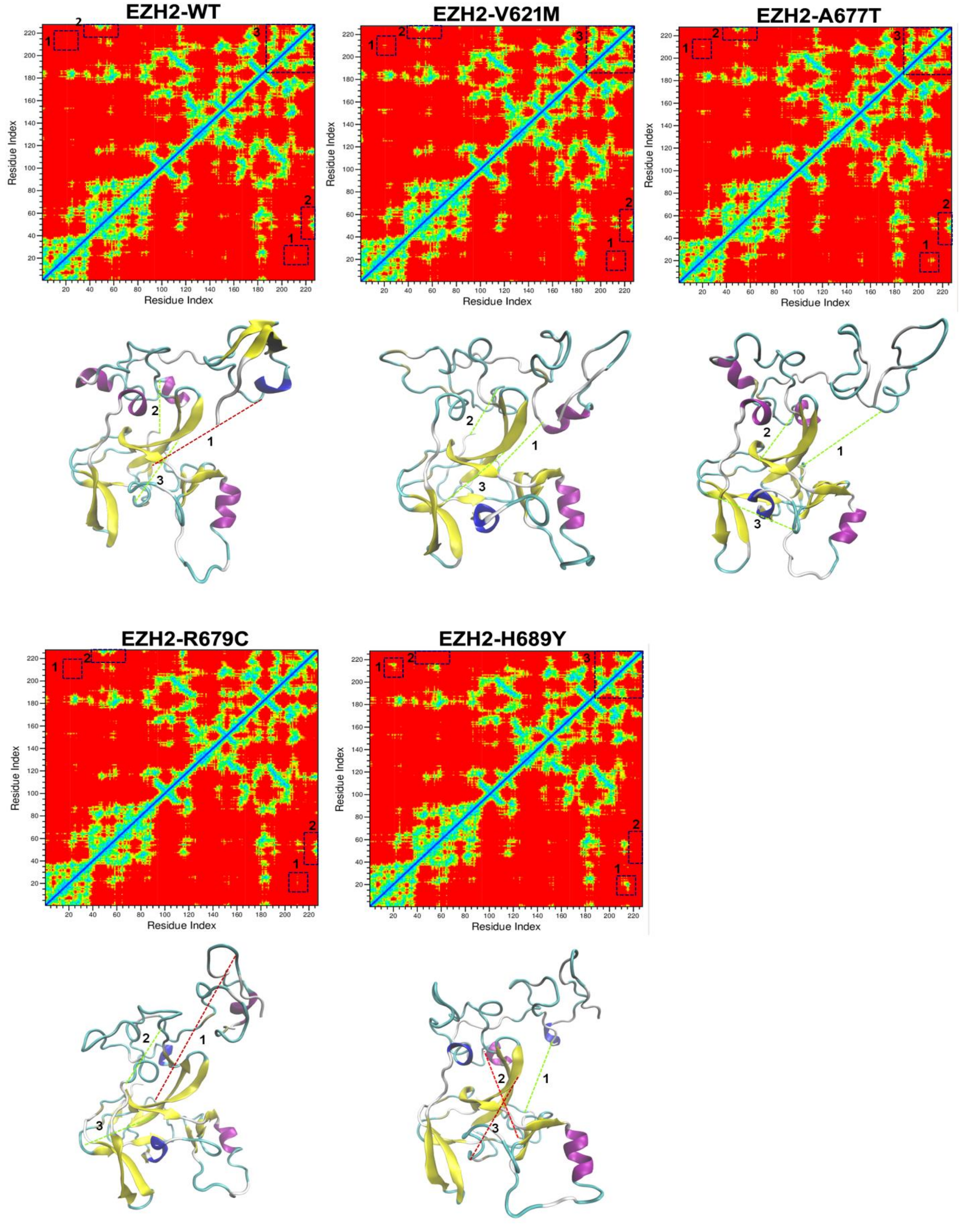
EZH2-WT
EZH2-V621M
EZH2-A677T
EZH2-R679C
EZH2-H689Y
Residue Index
Residue Index

### Figure S8

A

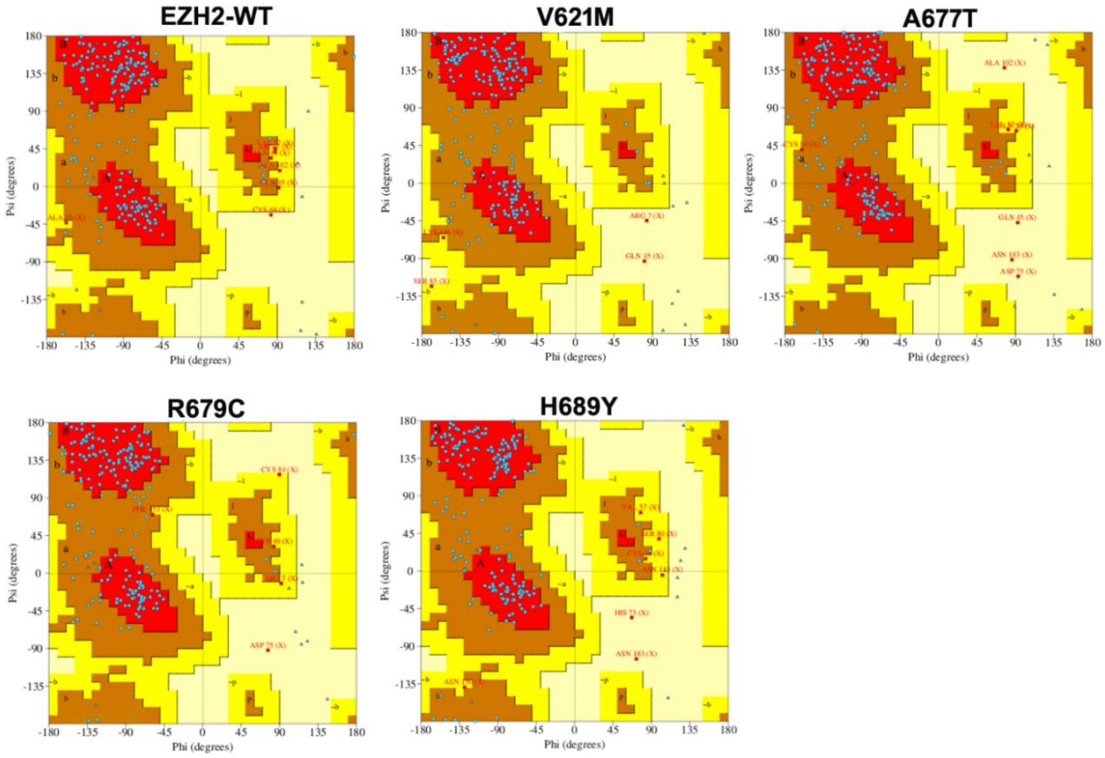

B

| Ramachandran Plot statistics | Calculated values | | | | |
|---|---|---|---|---|---|
| | EZH2-WT | V621M | A677T | R679C | H689Y |
| Residues in the most favoured regions [A,B,L] | 80.70 % | 79.70 % | 76.70 % | 75.7 % | 75.7 % |
| Residues in the additional allowed regions [a,b,l,p] | 15.80 % | 18.30 % | 19.80 % | 21.8 % | 20.8 % |
| Residues in the generously allowed regions [~a,~b,~l,~p] | 3.0 % | 1.0 % | 1.5 % | 2.0 % | 2.5 % |
| Residues in the disallowed regions [XX] | 0.5 % | 1.0 % | 2.0 % | 0.5 % | 1.0 % |

### Figure S9

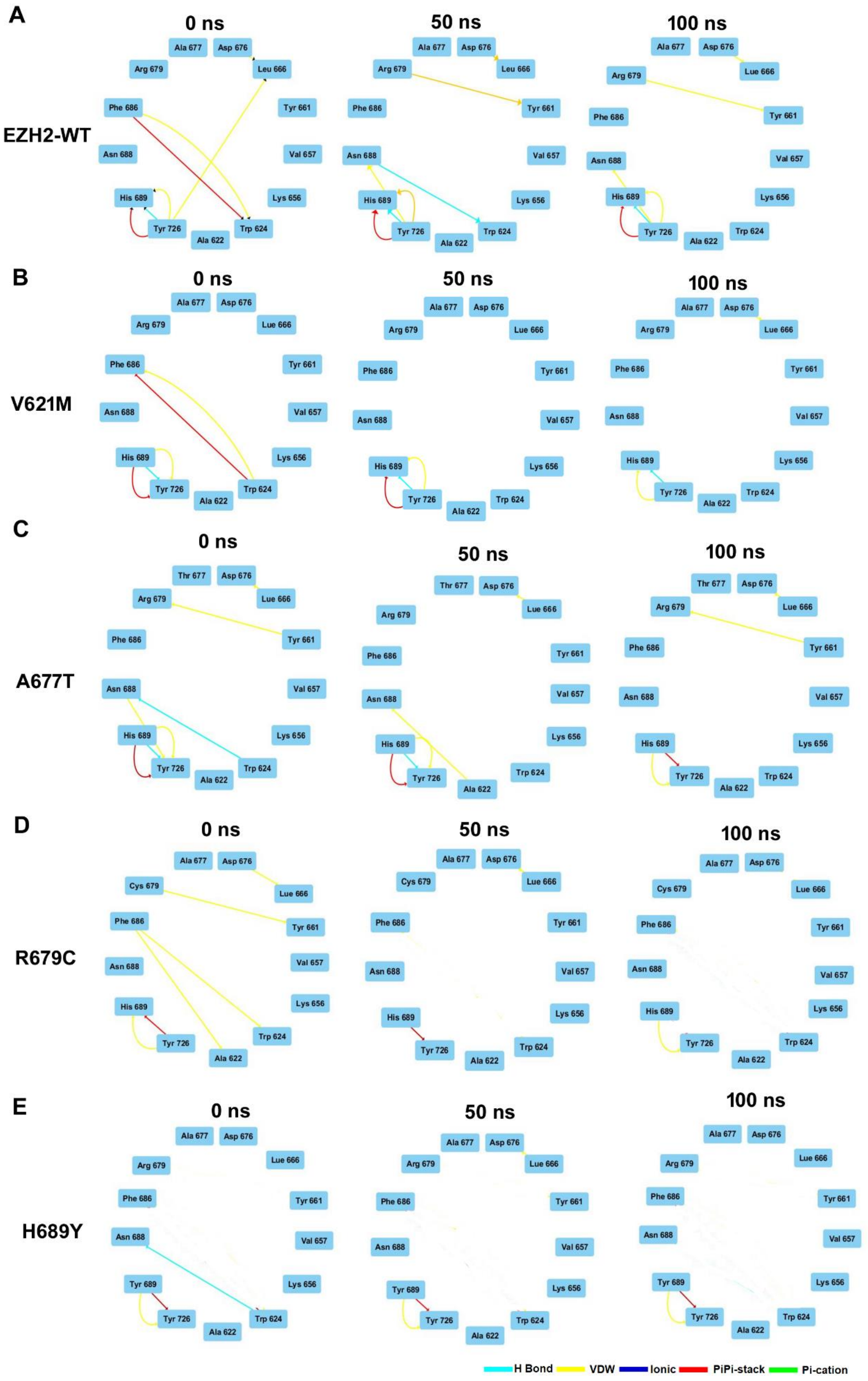

A
0 ns
50 ns
100 ns
EZH2-WT
B
V621M
C
A677T
D
R679C
E
H689Y
H Bond
VDW
Ionic
PiPi-stack
Pi-cation

### Figure S9

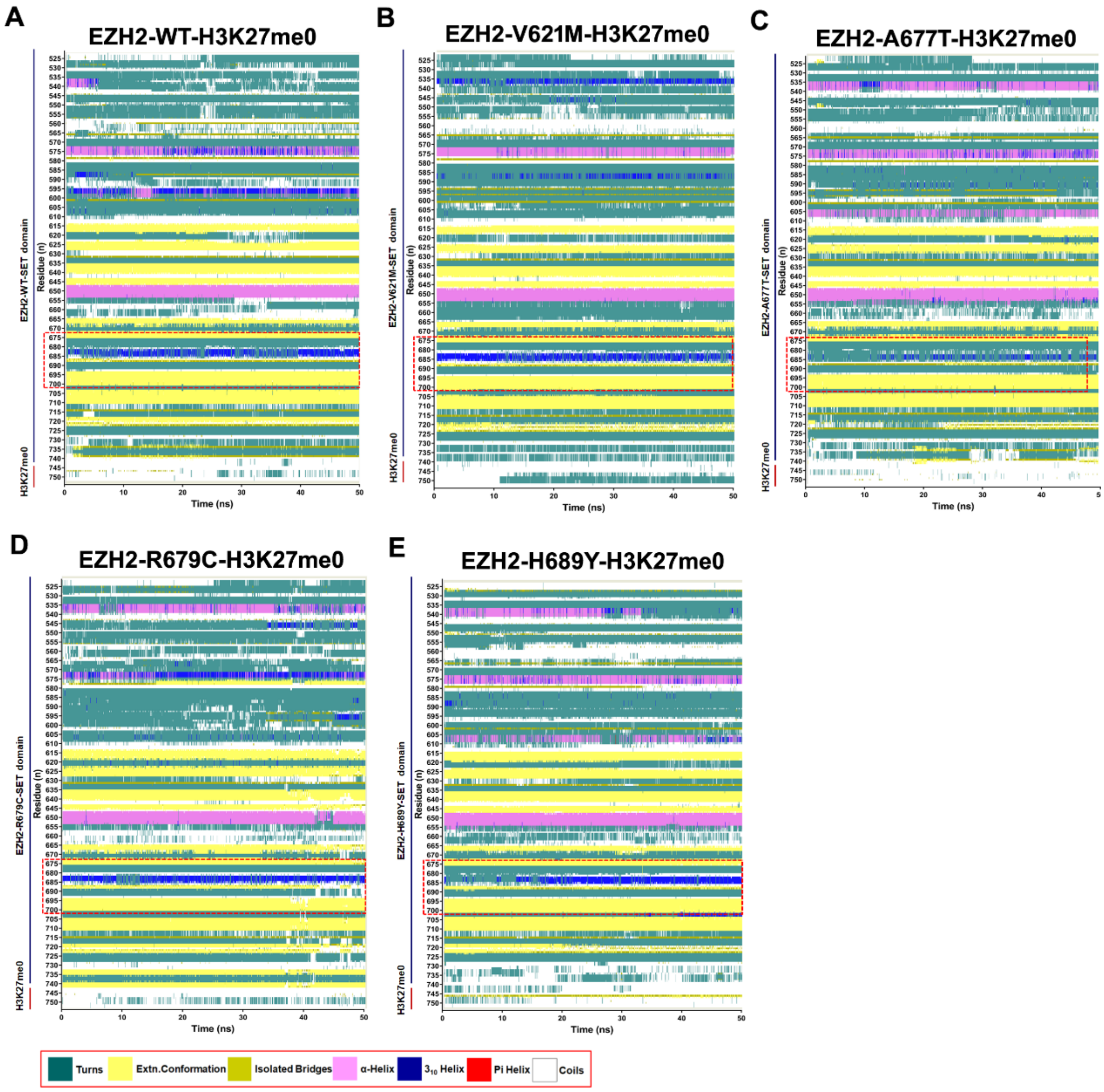
A
EZH2-WT-H3K27me0
B
EZH2-V621M-H3K27me0
C
EZH2-A677T-H3K27me0
D
EZH2-R679C-H3K27me0
E
EZH2-H689Y-H3K27me0
Residue (n)
Time (ns)
EZH2-WT-SET domain
EZH2-V621M-SET domain
EZH2-A677T-SET domain
EZH2-R679C-SET domain
EZH2-H689Y-SET domain
H3K27me0
Turns
Extn.Conformation
Isolated Bridges
α-Helix
3₁₀ Helix
Pi Helix
Coils

**Figure S10**

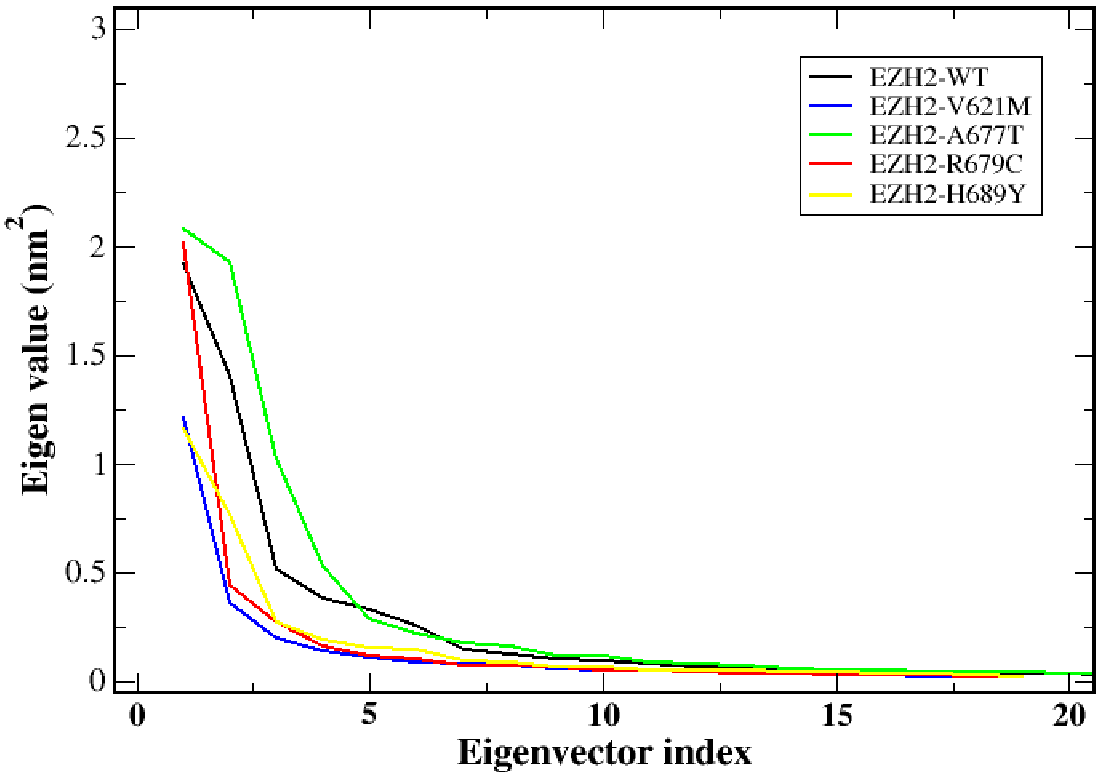
EZH2-WT
EZH2-V621M
EZH2-A677T
EZH2-R679C
EZH2-H689Y
Eigen value (nm$^2$)
3
2.5
2
1.5
1
0.5
0
0
5
10
15
20
Eigenvector index

### Figure S11

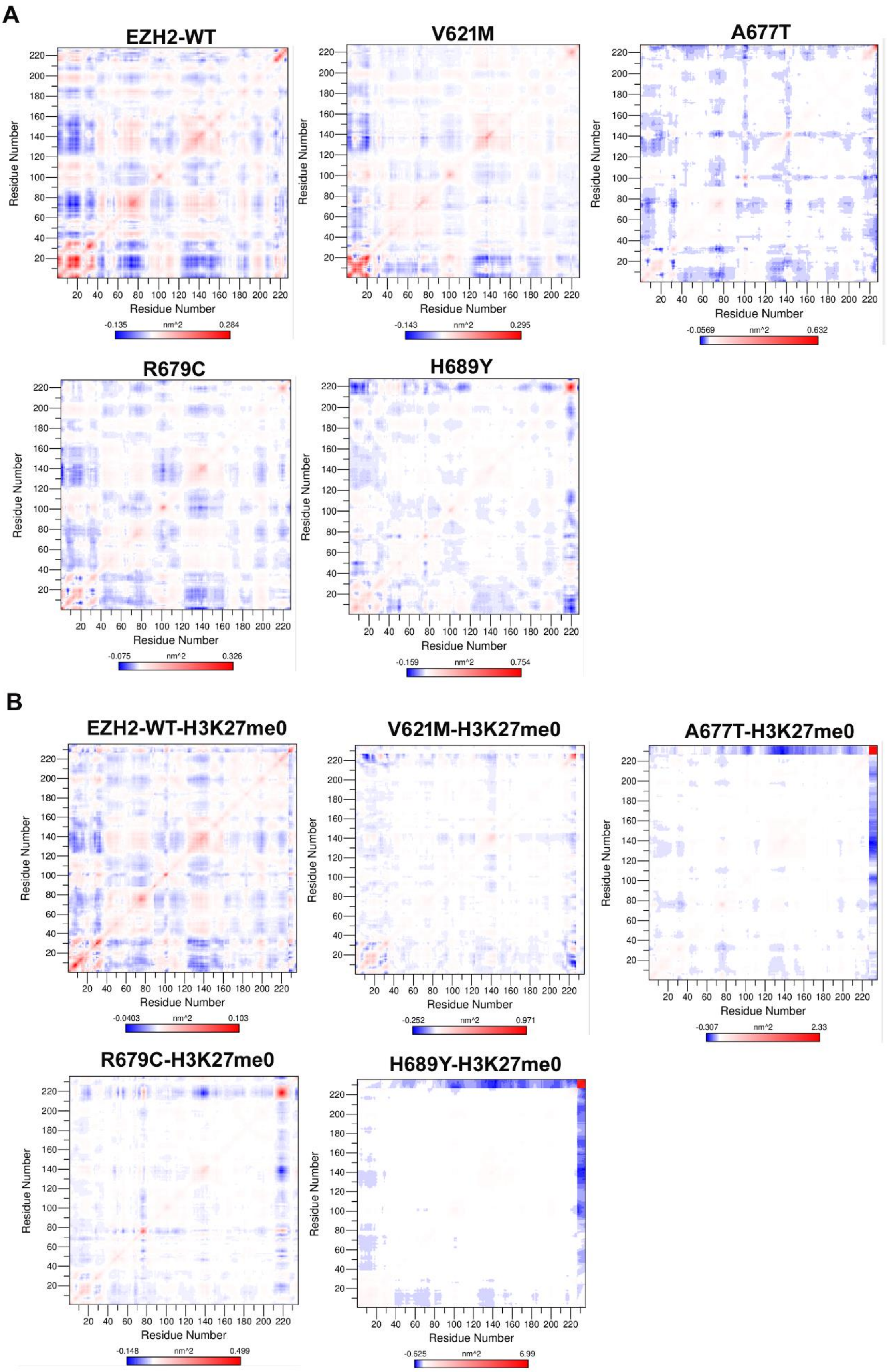
A
EZH2-WT
V621M
A677T
R679C
H689Y
Residue Number
nm^2
-0.135
0.284
-0.143
0.295
-0.0569
0.632
-0.075
0.326
-0.159
0.754
B
EZH2-WT-H3K27me0
V621M-H3K27me0
A677T-H3K27me0
R679C-H3K27me0
H689Y-H3K27me0
-0.0403
0.103
-0.252
0.971
-0.307
2.33
-0.148
0.499
-0.625
6.99

**Table S1.**

| SNP-ID | Nucleotide Change | Amino acid Change | Protein ID | PROVEAN score | Phenotype |
|---|---|---|---|---|---|
| rs52833659 | 553G>C | D185H | Q15910.2 | -2.458 | Neutral |
| rs151023145 | 965A>G | N317S | Q15910.2 | 1.132 | Neutral |
| rs193921147 | 2080C>T | H689Y | Q15910.2 | -5.425 | Deleterious |
| rs193921148 | 394C>T | P132S | Q15910.2 | -7.706 | Deleterious |
| rs199645805 | 165C>G | I55M | Q15910.2 | -0.28 | Neutral |
| rs201135441 | 1459G>A | A482T | Q15910.2 | -0.446 | Neutral |
| rs267601394 | 1937A>C | Y641S | Q15910.2 | -8.061 | Deleterious |
| rs267601395 | 1936T>A | Y641N | Q15910.2 | -8.029 | Deleterious |
| rs397515547 | 2044G>A | A677T | Q15910.2 | -3.753 | Deleterious |
| rs397515548 | 2233G>A | E740K | Q15910.2 | -3.467 | Deleterious |
| rs587778301 | 1844C>T | S610F | Q15910.2 | 0.533 | Neutral |
| rs587778302 | 1810G>A | V599M | Q15910.2 | -1.688 | Neutral |
| rs587778304 | 848C>T | T283M | Q15910.2 | -4.909 | Deleterious |
| rs587778305 | 1181G>C | G389A | Q15910.2 | -0.57 | Neutral |
| rs587783625 | 1876G>A | V621M | Q15910.2 | -2.522 | Deleterious |
| rs587783626 | 2050C>T | R679C | Q15910.2 | -7.172 | Deleterious |
| rs587783627 | 2236A>G | R741G | Q15910.2 | -5.401 | Deleterious |
| rs747009766 | 1555T>C | S514P | Q15910.2 | -4.235 | Deleterious |
| rs770006533 | 1474A>T | T487S | Q15910.2 | -2.723 | Deleterious |
| rs775407864 | 149T>C | L50S | Q15910.2 | -0.033 | Neutral |
| rs797044844 | 458A>G | Y153C | Q15910.2 | -8.974 | Deleterious |
| rs886039601 | 2007C>G | S664R | Q15910.2 | -4.706 | Deleterious |
| rs886062080 | 1383G>C | R456S | Q15910.2 | -1.765 | Neutral |
| rs1051056437 | 587G>C | R196T | Q15910.2 | -0.285 | Neutral |
| rs1057519833 | 2045C>G | A677G | Q15910.2 | -3.753 | Deleterious |
| rs1057519894 | 1922A>C | E636A | Q15910.2 | -5.571 | Deleterious |
| rs1057520183 | 745G>A | E249K | Q15910.2 | -2.578 | Deleterious |
| rs1060503430 | 1990G>T | D659Y | Q15910.2 | -8.47 | Deleterious |
| rs1064795225 | 2221T>C | Y736H | Q15910.2 | -3.837 | Deleterious |
| rs1064796617 | 550G>C | D184H | Q15910.2 | -0.929 | Neutral |
| rs1131692184 | 2213C>A | A733D | Q15910.2 | -1.389 | Neutral |
| rs6954744 | 958C>G | L315V | Q15910.2 | 0.21 | Neutral |
| rs61753264 | 506A>G | D168G | Q15910.2 | -6.791 | Deleterious |
| rs112029831 | 520C>T | R174C | NP_694543.1 | -0.916 | Neutral |
| rs112034331 | 1711T>C | S571P | NP_694543.1 | -1.581 | Neutral |
| rs139878257 | 1654G>A | A552T | NP_694543.1 | -0.469 | Neutral |
| rs141583753 | 232C>T | R78C | NP_694543.1 | -1.787 | Neutral |
| rs144316514 | 1003G>C | E33Q | NP_694543.1 | -1.235 | Neutral |
| rs147328633 | 1671T>G | S557R | NP_694543.1 | -1.586 | Neutral |
| rs189454324 | 268C>T | H90Y | NP_694543.1 | -5.173 | Deleterious |
| rs192731117 | 487G>A | D163N | NP_694543.1 | -0.647 | Neutral |
| rs200520401 | 631C>G | Q211E | NP_694543.1 | -0.847 | Neutral |
| rs200964386 | 923C>T | P308L | NP_694543.1 | -1.669 | Neutral |
| rs370444695 | 1199T>C | M400T | NP_694543.1 | -3.158 | Deleterious |

| | | | | | |
|---|---|---|---|---|---|
| rs372285596 | 647C>T | A216V | NP_694543.1 | -1.046 | Neutral |
| rs374699518 | 544G>C | D182H | NP_694543.1 | -2.666 | Deleterious |
| rs375168091 | 347G>A | G116E | NP_694543.1 | -7.687 | Deleterious |
| rs377467108 | 521G>T | R174L | NP_694543.1 | -0.275 | Neutral |
| rs537373788 | 542C>T | S181F | NP_694543.1 | -3.466 | Deleterious |
| rs553185801 | 1016C>T | T339I | NP_694543.1 | -0.189 | Neutral |
| rs561605379 | 1706G>A | R569Q | NP_694543.1 | -3.632 | Deleterious |
| rs566622851 | 1309A>C | I437L | NP_694543.1 | -0.307 | Neutral |
| rs568618347 | 649C>G | L217V | NP_694543.1 | -0.98 | Neutral |
| rs745554458 | 1521T>A | F507L | NP_694543.1 | -5.371 | Deleterious |
| rs746465165 | 428A>G | N143S | NP_694543.1 | 0.077 | Neutral |
| rs746749718 | 865T>G | L289V | NP_694543.1 | -1.794 | Neutral |
| rs746946161 | 178T>C | W60R | NP_694543.1 | -3.813 | Deleterious |
| rs747028969 | 530G>A | R177Q | NP_694543.1 | -0.564 | Neutral |
| rs747782211 | 890C>T | A297V | NP_694543.1 | -1.237 | Neutral |
| rs747933788 | 2041G>A | E681K | NP_694543.1 | -3.601 | Deleterious |
| rs748108870 | 298A>T | T100S | NP_004447.2 | -1.61 | Neutral |
| rs748458685 | 1238G>A | C413Y | NP_694543.1 | -10.169 | Deleterious |
| rs748860527 | 902C>T | A301V | NP_694543.1 | -0.461 | Neutral |
| rs748915411 | 1084A>C | K362Q | NP_694543.1 | -0.91 | Neutral |
| rs749239554 | 1468C>T | P490S | NP_694543.1 | -4.527 | Deleterious |
| rs749498698 | 473A>G | E158G | NP_694543.1 | -1.352 | Neutral |
| rs749847544 | 1743C>G | D581E | NP_694543.1 | -3.534 | Deleterious |
| rs751723382 | 1767C>G | I589M | NP_694543.1 | -1.226 | Neutral |
| rs752284693 | 155G>A | R52L | NP_694543.1 | -1.567 | Neutral |
| rs753739962 | 1789G>C | E597Q | NP_694543.1 | -2.788 | Deleterious |
| rs754291699 | 1606C>G | L536V | NP_694543.1 | -2.926 | Deleterious |
| rs754403133 | 118A>G | S40G | NP_694543.1 | -1.751 | Neutral |
| rs755823072 | 590C>T | T197I | NP_694543.1 | -3.983 | Deleterious |
| rs756942104 | 673A>G | I225V | NP_694543.1 | -0.81 | Neutral |
| rs757534855 | 1765A>T | I589F | NP_694543.1 | -2.768 | Deleterious |
| rs757601923 | 959C>T | P320L | NP_694543.1 | -1.51 | Neutral |
| rs757807865 | 719A>C | H240P | NP_694543.1 | -9.874 | Deleterious |
| rs758449513 | 491T>G | L164R | NP_694543.1 | -0.227 | Neutral |
| rs759313335 | 379A>T | I127L | NP_694543.1 | -0.822 | Neutral |
| rs759576587 | 274C>T | P92S | NP_004447.2 | -1.118 | Neutral |
| rs761247972 | 131C>A | S44Y | NP_694543.1 | -2.201 | Neutral |
| rs761656628 | 1859A>G | D620G | NP_694543.1 | -6.593 | Deleterious |
| rs761834990 | 1639C>T | L547F | NP_694543.1 | -1.494 | Neutral |
| rs763225504 | 77A>G | H26R | XP_011514188.1 | 0.656 | Neutral |
| rs763682209 | 700C>A | Q234K | NP_694543.1 | -1.285 | Neutral |
| rs764649740 | 30G>C | L10N | NP_004447.2 | -1.746 | Neutral |
| rs765147666 | 503G>A | R168Q | NP_694543.1 | -0.451 | Neutral |
| rs765323739 | 269A>T | D90V | NP_004447.2 | -2.628 | Deleterious |
| rs765380980 | 255T>A | D85E | NP_694543.1 | -3.838 | Deleterious |
| rs765768601 | 1304C>A | S435Y | NP_694543.1 | -2.408 | Neutral |
| rs765980265 | 241A>G | R81G | NP_694543.1 | -1.726 | Neutral |
| rs766387427 | 1813G>A | E605K | NP_694543.1 | -3.768 | Deleterious |
| rs766447146 | 295G>A | V99I | NP_694543.1 | -0.538 | Neutral |

| | | | | | |
|---|---|---|---|---|---|
| rs766875832 | 800A>C | Y267S | NP_694543.1 | -1.529 | Neutral |
| rs766928732 | 121A>T | M41L | NP_694543.1 | -0.261 | Neutral |
| rs767489605 | 489T>G | D163E | NP_694543.1 | 0.137 | Neutral |
| rs767528515 | 1633C>T | L545F | NP_694543.1 | -3.768 | Deleterious |
| rs768512235 | 1467G>T | Q489H | NP_694543.1 | -1.802 | Neutral |
| rs768812143 | 158C>T | T53M | NP_694543.1 | -1.311 | Neutral |
| rs769298548 | 1091A>G | E364G | NP_694543.1 | -0.871 | Neutral |
| rs770030757 | 510T>A | D170E | NP_694543.1 | -0.991 | Neutral |
| rs770327434 | 1154T>C | I385T | NP_694543.1 | -1.013 | Neutral |
| rs770380532 | 423A>T | Q141H | NP_694543.1 | -2.733 | Deleterious |
| rs771139896 | 529C>T | R177W | NP_694543.1 | -2.101 | Neutral |
| rs771352080 | 52C>T | R18C | NP_694543.1 | -4.988 | Deleterious |
| rs771467281 | 1978G>A | V660I | NP_694543.1 | -0.903 | Neutral |
| rs771888727 | 169C>T | L57F | XP_011514188.1 | -0.224 | Neutral |
| rs772530358 | 148T>G | L50V | XP_011514188.1 | -0.371 | Neutral |
| rs773141417 | 278C>G | T93R | XP_011514188.1 | -4.291 | Deleterious |
| rs773492931 | 820A>G | T274A | XP_011514188.1 | -0.362 | Neutral |
| rs774270705 | 1067A>C | K356T | XP_011514188.1 | -0.685 | Neutral |
| rs775295296 | 1126C>G | Q376E | XP_011514188.1 | -1.043 | Neutral |
| rs775942317 | 929G>C | R310P | XP_011514188.1 | -1.461 | Neutral |
| rs776246781 | 1150A>C | N384H | XP_011514188.1 | -1.29 | Neutral |
| rs776925814 | 1417G>A | G473S | XP_011514188.1 | 1.649 | Neutral |
| rs776984937 | 49A>C | K17Q | XP_011514188.1 | -2.431 | Neutral |
| rs778968366 | 457G>A | D153N | XP_011514188.1 | -0.739 | Neutral |
| rs779136188 | 1243A>C | I415L | XP_011514188.1 | -0.988 | Neutral |
| rs779629814 | 2118C>G | I706M | XP_011514188.1 | -0.599 | Neutral |
| rs779757594 | 1088A>G | D363G | XP_011514188.1 | -1.296 | Neutral |
| rs779779653 | 1531C>A | L511I | XP_011514201.1 | -1.447 | Neutral |
| rs779951996 | 571A>G | M191V | NP_694543.1 | -1.643 | Neutral |
| rs780251816 | 476A>C | E159A | NP_694543.1 | -1.851 | Neutral |
| rs780959381 | 670A>G | N224D | NP_694543.1 | -4.306 | Deleterious |
| rs781275057 | 1954G>C | V652L | NP_694543.1 | -1.485 | Neutral |
| rs781407066 | 1759A>G | I587V | NP_694543.1 | -0.678 | Neutral |
| rs781431240 | 955C>T | L319F | NP_694543.1 | -0.162 | Neutral |
| rs781468426 | 953G>A | R318Q | NP_694543.1 | -0.456 | Neutral |
| rs866144035 | 1226A>G | Y409C | NP_694543.1 | -3.413 | Deleterious |
| rs867587253 | 449A>G | D150G | NP_694543.1 | -1.965 | Neutral |
| rs868477296 | 1294G>T | V432F | NP_694543.1 | -1.892 | Neutral |
| rs886735712 | 38T>C | V13A | XP_024302448.1 | -0.059 | Neutral |
| rs886990256 | 2079T>A | D693E | NP_694543.1 | 0 | Neutral |
| rs897492574 | 1232A>G | N411S | NP_694543.1 | -4.485 | Deleterious |
| rs897885462 | 899G>A | C300Y | NP_004447.2 | 0.029 | Neutral |
| rs898443241 | 25G>T | A9S | XP_024302448.1 | 0.625 | Neutral |
| rs915879954 | 7T>C | Y3H | XP_011514185.1 | 0.246 | Neutral |
| rs917585556 | 551T>C | I184T | NP_694543.1 | -4.313 | Deleterious |
| rs938955302 | 98A>C | N33T | XP_011514188.1 | -1.979 | Neutral |
| rs939117179 | 49A>T | T17S | XP_011514193.1 | 0.564 | Neutral |
| rs939686972 | 35C>G | P12Q | XP_011514193.1 | 0.802 | Neutral |
| rs961766373 | 70A>G | I24V | XP_011514188.1 | -0.056 | Neutral |

| | | | | | |
|---|---|---|---|---|---|
| rs968299371 | 310G>A | V104I | NP_004447.2 | -0.741 | Neutral |
| rs972308238 | 139C>G | Q47G | NP_694543.1 | -3.077 | Deleterious |
| rs973062748 | 524C>T | P174L | NP_694543.1 | -0.275 | Neutral |
| rs979372067 | 1626C>A | D542E | NP_694543.1 | -3.768 | Deleterious |
| rs980295835 | 157C>T | L53F | XP_011514188.1 | -0.642 | Neutral |
| rs982908069 | 595G>A | E199L | NP_694543.1 | -6.261 | Deleterious |
| rs987432794 | 130T>G | S44A | XP_011514193.1 | 0.113 | Neutral |
| rs987568723 | 1945A>C | N649H | NP_694543.1 | -4.524 | Deleterious |
| rs988151352 | 1469C>T | P490L | NP_694543.1 | -6.095 | Deleterious |
| rs990847755 | 2020A>G | K674E | NP_694543.1 | -3.335 | Deleterious |
| rs997068128 | 14T>G | L5R | XP_024302448.1 | 2.189 | Neutral |
| rs999855448 | 305A>G | N102S | NP_004447.2 | -2.764 | Deleterious |
| rs1005840534 | 625A>G | T209A | NP_694543.1 | -3.206 | Deleterious |
| rs1015647802 | 122G>C | S41T | XP_011514188.1 | 0.521 | Neutral |
| rs1022187456 | 221A>T | D74V | XP_011514188.1 | -0.055 | Neutral |
| rs1022373846 | 205C>T | H69Y | NP_694543.1 | -2.007 | Neutral |
| rs1029480467 | 143C>T | T48M | XP_011514193.1 | -0.224 | Neutral |
| rs1035066688 | 8A>T | Q3L | NP_694543.1 | -0.977 | Neutral |
| rs1048763359 | 1109A>G | E370G | NP_694543.1 | -2.2 | Neutral |
| rs1054582039 | 1298A>C | K433T | NP_694543.1 | -3.829 | Deleterious |
| rs1158237595 | 53G>A | R18Q | NP_694543.1 | -2.419 | Neutral |
| rs1158852320 | 944G>A | R315K | NP_694543.1 | -0.296 | Neutral |
| rs1163934928 | 1151A>G | N384S | NP_694543.1 | -0.856 | Neutral |
| rs1164182968 | 626C>T | T209I | NP_694543.1 | -3.258 | Deleterious |
| rs1164610428 | 1550G>A | R517H | NP_694543.1 | -4.536 | Deleterious |
| rs1167346838 | 83C>G | A28G | XP_011514193.1 | -0.389 | Neutral |
| rs1168301922 | 1324C>G | P442A | NP_694543.1 | -2.505 | Deleterious |
| rs1172023956 | 142A>C | T48P | XP_011514193.1 | -0.562 | Neutral |
| rs1177540949 | 8T>C | I3T | XP_011514193.1 | 1.597 | Neutral |
| rs1179661739 | 495G>C | E165D | NP_694543.1 | -0.776 | Neutral |
| rs1180219356 | 265T>A | L265M | NP_004447.2 | -8.503 | Deleterious |
| rs1181217229 | 72T>G | S24R | XP_011514193.1 | 0.675 | Neutral |
| rs1182158215 | 932C>T | P311L | NP_694543.1 | -0.94 | Neutral |
| rs1182369323 | 251C>T | S84L | NP_004447.2 | -2.111 | Neutral |
| rs1182707339 | 79C>T | P27S | XP_011514193.1 | -0.094 | Neutral |
| rs1186719861 | 2T>G | M1R | XP_011514193.1 | 1.51 | Neutral |
| rs1189608375 | 782A>T | H261L | NP_694543.1 | -6.32 | Deleterious |
| rs1189939128 | 1400T>C | I467T | NP_694543.1 | -4.523 | Deleterious |
| rs1190643983 | 103C>A | P35T | XP_016867307.1 | 1.445 | Neutral |
| rs1191492481 | 1064A>T | D355V | NP_694543.1 | -1.69 | Neutral |
| rs1193491117 | 952C>T | R318W | NP_694543.1 | -2.013 | Neutral |
| rs1196167123 | 67G>A | G23S | XP_024302448.1 | 0.604 | Neutral |
| rs1196167328 | 98G>A | R33K | NP_694543.1 | 1.273 | Neutral |
| rs1198807049 | 1214T>C | I405T | NP_694543.1 | 0.449 | Neutral |
| rs1199671048 | 136C>T | R46C | NP_694543.1 | -3.796 | Deleterious |
| rs1200905474 | 1046C>T | T349M | NP_694543.1 | -1.238 | Neutral |
| rs1203188103 | 1510T>C | S518P | NP_001190177.1 | 1.345 | Neutral |
| rs1209325621 | 100C>T | R34C | XP_011514193.1 | -0.227 | Neutral |
| rs1209538167 | 118C>G | L40V | XP_011514188.1 | -0.034 | Neutral |

| | | | | | |
|---|---|---|---|---|---|
| rs1211566129 | 59T>G | L20R | XP_011514188.1 | 0.427 | Neutral |
| rs1212616016 | 583A>G | K195E | NP_694543.1 | -3.077 | Deleterious |
| rs1214137197 | 964A>G | N322D | NP_694543.1 | 0.417 | Neutral |
| rs1215160919 | 65A>C | E22A | NP_694543.1 | -4.859 | Deleterious |
| rs1218303603 | 233G>A | R78H | NP_694543.1 | -0.47 | Neutral |
| rs1220875602 | 655C>T | P219S | NP_694543.1 | -4.77 | Deleterious |
| rs1224767362 | 634C>G | Q212E | NP_694543.1 | -0.907 | Neutral |
| rs1225047502 | 1255A>C | I419L | NP_694543.1 | -0.457 | Neutral |
| rs1225799902 | 19C>T | K7C | NP_694543.1 | -1.932 | Neutral |
| rs1228980656 | 289C>T | P97S | NP_004447.2 | -3.929 | Deleterious |
| rs1229068307 | 1428C>G | N520L | NP_004447.2 | -6.032 | Deleterious |
| rs1233462075 | 1200G>A | M400I | NP_694543.1 | -0.278 | Neutral |
| rs1233687730 | 650T>C | L217P | NP_694543.1 | -3.232 | Deleterious |
| rs1234137926 | 70A>G | S24G | XP_011514193.1 | -1.095 | Neutral |
| rs1234158247 | 412G>A | A138T | NP_694543.1 | -2.662 | Deleterious |
| rs1236254798 | 40T>C | F14L | XP_011514188.1 | 0.544 | Neutral |
| rs1238469391 | 1A>T | M1L | XP_011514185.1 | 0.02 | Neutral |
| rs1244095507 | 131C>T | S44L | XP_011514193.1 | 1.096 | Neutral |
| rs1253076713 | 46G>A | D16N | XP_024302448.1 | 1.356 | Neutral |
| rs1253081994 | 902A>G | N301S | NP_004447.2 | 0.225 | Neutral |
| rs1254820988 | 1066A>G | K356E | NP_694543.1 | -0.933 | Neutral |
| rs1259698211 | 37G>A | D13N | XP_011514188.1 | 0.029 | Neutral |
| rs1260470042 | 134A>G | N45S | NP_694543.1 | -4.257 | Deleterious |
| rs1262192022 | 128A>G | E43G | XP_011514193.1 | -0.395 | Neutral |
| rs1263098237 | 439G>A | D147N | NP_694543.1 | -0.964 | Neutral |
| rs1263153853 | 1373G>A | R458Q | NP_694543.1 | -3.328 | Deleterious |
| rs1263788302 | 519C>G | S173R | NP_694543.1 | -1.287 | Neutral |
| rs1265059186 | 996T>G | N332K | NP_694543.1 | -0.57 | Neutral |
| rs1266279264 | 5C>T | A2V | XP_024302448.1 | -0.383 | Neutral |
| rs1268069858 | 1681C>T | R561C | NP_004447.2 | -7.285 | Deleterious |
| rs1273062625 | 242G>C | R81T | NP_694543.1 | -0.823 | Neutral |
| rs1280677426 | 130G>C | V44L | XP_011514188.1 | 0.572 | Neutral |
| rs1281404133 | 997G>A | V333M | NP_694543.1 | -0.127 | Neutral |
| rs1283942043 | 107G>A | G36D | XP_011514193.1 | 1.325 | Neutral |
| rs1284186657 | 95C>T | A32V | XP_011514193.1 | 0.558 | Neutral |
| rs1285260202 | 962A>C | N321T | NP_694543.1 | -0.898 | Neutral |
| rs1291033053 | 110A>C | E37A | NP_694543.1 | -4.47 | Deleterious |
| rs1292037761 | 1048G>C | G350R | NP_694543.1 | -1.352 | Neutral |
| rs1298174513 | 1009A>C | K337Q | NP_694543.1 | -1.099 | Neutral |
| rs1299959949 | 247G>T | V83L | NP_694543.1 | -2.845 | Deleteriou |
| rs1305651544 | 22T>G | L8V | XP_011514193.1 | 0.103 | Neutral |
| rs1308485787 | 446A>C | D149A | NP_694543.1 | -0.336 | Neutral |
| rs1309469934 | 898A>G | T300A | NP_694543.1 | 1.155 | Neutral |
| rs1315592469 | 804G>C | K268N | NP_694543.1 | -3.986 | Deleterious |
| rs1315928462 | 1135A>G | I379V | NP_694543.1 | -0.02 | Neutral |
| rs1318022205 | 1475A>G | D492G | NP_694543.1 | -6.618 | Deleterious |
| rs1319998961 | 280C>G | Q94E | NP_004447.2 | -2.23 | Neutral |
| rs1320364194 | 64G>A | E22L | XP_011514193.1 | 0.085 | Neutral |
| rs1321951994 | 1903G>A | V635M | NP_694543.1 | -2.583 | Deleterious |

| | | | | | |
|---|---|---|---|---|---|
| rs1322202663 | 442G>C | D148H | NP_694543.1 | -2.57 | Deleterious |
| rs1323501058 | 16C>T | P6S | XP_011514193.1 | 1.552 | Neutral |
| rs1324786734 | 343C>G | P115A | NP_004447.2 | -6.311 | Deleterious |
| rs1326457692 | 64T>G | S22A | XP_011514188.1 | 0.03 | Neutral |
| rs1328717365 | 1102T>G | S368A | NP_694543.1 | -1.216 | Neutral |
| rs1330328545 | 58C>T | L20F | XP_011514188.1 | -0.033 | Neutral |
| rs1330757055 | 983C>G | T328S | NP_694543.1 | -0.555 | Neutral |
| rs1333628386 | 137G>A | R46H | NP_694543.1 | -2.468 | Neutral |
| rs1334047435 | 112G>A | A38T | XP_011514193.1 | 0.153 | Neutral |
| rs1336390351 | 1292G>C | R431T | NP_694543.1 | 0.096 | Neutral |
| rs1336750860 | 200G>A | R67H | XP_011514188.1 | 0.383 | Neutral |
| rs1338552132 | 148T>A | Y50N | XP_011514188.1 | 0.157 | Neutral |
| rs1338606877 | 50C>G | T17S | XP_011514193.1 | 0.564 | Neutral |
| rs1340007682 | 1366A>C | D556Q | NP_694543.1 | -1.976 | Neutral |
| rs1342246023 | 496G>T | D166Y | NP_694543.1 | -3.131 | Deleterious |
| rs1345461419 | 1898A>T | D633V | NP_694543.1 | -7.948 | Deleterious |
| rs1346782884 | 440A>G | D147G | NP_694543.1 | -1.436 | Neutral |
| rs1348759539 | 1421C>T | S474F | NP_694543.1 | -4.771 | Deleterious |
| rs1352634479 | 965A>C | N322T | NP_694543.1 | -0.804 | Neutral |
| rs1354940290 | 667C>T | P223S | NP_694543.1 | -7.631 | Deleterious |
| rs1355148527 | 1078G>C | E360Q | NP_694543.1 | -0.564 | Neutral |
| rs1356354491 | 88G>C | A30P | XP_011514193.1 | 0.362 | Neutral |
| rs1359374351 | 9G>C | Q3H | NP_694543.1 | -0.898 | Neutral |
| rs1359796591 | 1283A>G | Y428C | NP_694543.1 | -7.772 | Deleterious |
| rs1362588674 | 47A>T | H16L | XP_011514188.1 | 0.342 | Neutral |
| rs1365724658 | 60C>G | D20E | XP_011514193.1 | 0.566 | Neutral |
| rs1366762452 | 919C>G | P307A | NP_694543.1 | -0.542 | Neutral |
| rs1369190067 | 163A>G | I55V | NP_694543.1 | -0.115 | Neutral |
| rs1371460362 | 32G>C | G11A | XP_011514193.1 | 0.746 | Neutral |
| rs1371803981 | 58G>A | D20N | XP_011514193.1 | 0.131 | Neutral |
| rs1373185092 | 1349C>T | P450L | NP_694543.1 | -8.492 | Deleterious |
| rs1375274910 | 2034G>C | Q678H | NP_694543.1 | -3.611 | Deleterious |
| rs1375591963 | 1040C>T | T34I | NP_694543.1 | -5.955 | Deleterious |
| rs1377401772 | 98C>T | A33V | XP_011514193.1 | 0.918 | Neutral |
| rs1378812518 | 935G>C | G312A | NP_694543.1 | -1.083 | Neutral |
| rs1382627046 | 494A>G | E165G | NP_694543.1 | -2.761 | Deleterious |
| rs1383600888 | 1661A>G | H554R | NP_694543.1 | -2.099 | Neutral |
| rs1387683954 | 1020C>A | D340E | NP_694543.1 | -1.482 | Neutral |
| rs1388815758 | 235G>A | G79R | NP_694543.1 | 0.637 | Neutral |
| rs1389657572 | 151G>A | E51K | NP_694543.1 | -0.76 | Neutral |
| rs1390180981 | 901G>A | A301T | NP_694543.1 | -0.898 | Neutral |
| rs1394570516 | 882G>C | G294D | NP_694543.1 | -1.977 | Neutral |
| rs1395172893 | 10G>A | G4R | XP_011514193.1 | 2.293 | Neutral |
| rs1395843139 | 1986G>A | M662I | NP_694543.1 | -3.602 | Deleterious |
| rs1396484737 | 1999C>A | H667N | NP_694543.1 | -6.069 | Deleterious |
| rs1400051032 | 32G>A | G11E | NP_694543.1 | -0.267 | Neutral |
| rs1402189141 | 153A>T | E51D | NP_694543.1 | -1.167 | Neutral |
| rs1405534761 | 179C>T | P60L | XP_011514188.1 | -1.022 | Neutral |
| rs1406929961 | 52C>T | R18W | XP_011514193.1 | 0.013 | Neutral |

| | | | | | |
|---|---|---|---|---|---|
| rs1410650509 | 110A>G | D37G | XP_011514193.1 | -0.764 | Neutral |
| rs1411356297 | 80C>G | P27R | XP_011514193.1 | 1.463 | Neutral |
| rs1413310167 | 134A>G | Y45C | XP_011514188.1 | 0.323 | Neutral |
| rs1414257152 | 178C>G | P60A | XP_011514188.1 | -1.202 | Neutral |
| rs1417188418 | 116G>A | R39Q | XP_011514193.1 | 0.278 | Neutral |
| rs1418025724 | 1331A>C | E444A | NP_694543.1 | -2.551 | Deleterious |
| rs1418949011 | 13C>G | L5V | XP_011514193.1 | 0.362 | Neutral |
| rs1423687581 | 80A>C | N27T | XP_011514188.1 | -0.472 | Neutral |
| rs1425631167 | 1198A>G | M400V | NP_694543.1 | -1.136 | Neutral |
| rs1426017767 | 427A>G | N143D | NP_694543.1 | -0.72 | Neutral |
| rs1426509356 | 1683A>C | Q561H | XP_011514185.1 | -1.87 | Neutral |
| rs1427041861 | 907C>T | R303W | NP_694543.1 | -2.342 | Neutral |
| rs1428739066 | 2117T>G | I706S | NP_694543.1 | -2.804 | Deleterious |
| rs1431254558 | 1286A>G | E429G | NP_694543.1 | -3.976 | Deleterious |
| rs1434330368 | 292T>A | L98I | NP_004447.2 | 0.118 | Neutral |
| rs1437977406 | 1955T>C | V652A | NP_694543.1 | -3.319 | Deleterious |
| rs1439519518 | 31G>C | G11R | XP_011514193.1 | 2.312 | Neutral |
| rs1439884659 | 11G>C | G4A | XP_011514193.1 | 0.667 | Neutral |
| rs1440638451 | 397G>C | V133L | NP_694543.1 | -1.827 | Neutral |
| rs1441187386 | 940C>T | R314C | NP_694543.1 | -2.272 | Neutral |
| rs1441305206 | 994A>C | N334H | NP_694543.1 | -0.91 | Neutral |
| rs1443131834 | 1574C>T | A525V | NP_694543.1 | -3.368 | Deleterious |
| rs1443557345 | 92C>T | A31V | XP_011514193.1 | 0.301 | Neutral |
| rs1446181129 | 59A>T | D20V | XP_011514193.1 | 0.448 | Neutral |
| rs1446361012 | 332T>A | I111K | NP_694543.1 | -6.559 | Deleterious |
| rs1451233310 | 41C>T | A14V | XP_011514193.1 | 0.403 | Neutral |
| rs1451278890 | 1993G>A | G665S | NP_694543.1 | -5.173 | Deleterious |
| rs1454209655 | 22T>C | S8P | NP_694543.1 | -1.2 | Neutral |
| rs1454562282 | 1121G>A | R374Q | NP_694543.1 | -0.233 | Neutral |
| rs1456525510 | 55T>C | W19R | XP_011514193.1 | -1.236 | Neutral |
| rs1456751008 | 431A>G | D144G | NP_694543.1 | -2.089 | Neutral |
| rs1459861646 | 1583A>C | N528T | NP_694543.1 | -5.517 | Deleterious |
| rs1462379572 | 1372C>T | R458T | NP_694543.1 | -5.408 | Deleterious |
| rs1465177913 | 1031A>G | E344G | NP_694543.1 | -1.774 | Neutral |
| rs1465319325 | 86C>T | A29V | XP_011514193.1 | 1.335 | Neutral |
| rs1466183869 | 401A>C | G134A | NP_694543.1 | -5.572 | Deleterious |
| rs1468342429 | 10G>A | A4T | XP_011514188.1 | -0.511 | Neutral |
| rs1468914332 | 1057A>G | N353D | NP_694543.1 | -0.485 | Neutral |
| rs1473228269 | 1429C>T | H477Y | NP_694543.1 | -4.317 | Deleterious |
| rs1474211340 | 1277A>C | H426P | XP_011514203.1 | 1.382 | Neutral |
| rs1475241273 | 206A>G | H69R | NP_694543.1 | -0.297 | Neutral |
| rs1475881563 | 139G>C | G47R | XP_011514193.1 | 1.291 | Neutral |
| rs1476603937 | 1751G>C | G584A | NP_694543.1 | -4.98 | Deleterious |
| rs1478572656 | 94T>G | W32G | XP_011514188.1 | -1.184 | Neutral |
| rs1479961823 | 116G>C | S39T | XP_016867307.1 | 0.008 | Neutral |
| rs1480044679 | 1970A>G | Y657C | NP_694543.1 | -7.79 | Deleterious |
| rs1481097067 | 1153A>G | I385V | NP_694543.1 | 0.016 | Neutral |
| rs1483639186 | 1694G>A | C565Y | NP_694543.1 | -2.628 | Deleterious |
| rs1484428746 | 290A>T | D97V | NP_694543.1 | -8.114 | Deleterious |

| | | | | | |
|---|---|---|---|---|---|
| rs1484617559 | 121A>G | S41G | XP_011514188.1 | -0.813 | Neutral |
| rs1488148920 | 26A>T | E9V | NP_694543.1 | -1.68 | Neutral |
| rs1488870104 | 358G>A | G120R | NP_694543.1 | -6.951 | Deleterious |
| rs1488978716 | 109G>A | A37T | XP_011514188.1 | 0.068 | Neutral |

**Supplementary Figures and Table Legends:**

**Figure S1.** Multiple sequence alignment of significant sequences of EZH2 with their closely related sequence processed through MUSCLE program.

**Figure S2**. Evolutionary analyses were conducted in MEGA-X and visualize midpoint rooted tree in FigTree v1.4.4. The evolutionary history was inferred using the Neighbor-Joining method. The bootstrap consensus tree inferred from 1000 replicates is taken to represent the evolutionary history of the taxa analyzed. The evolutionary distances were computed using the p-distance method and are in the units of the number of amino acid differences per site. The rate variation among sites was modeled with a gamma distribution (shape parameter = 1). All positions containing gaps and missing data were eliminated (complete deletion). The color of the tips blue and black indicates significant and non-significant sequences, respectively.

**Figure S3.** Distance matrices showing the smallest distance the residue pair in EZH2-WT (A), EZH2-V621, EZH2-A677T, EZH2-R679C and EZH2-H689Y with their respective conformational poses. The circles were used to differentiate 1, 2 and 3 changes in the mutants when compare to wild-type.

**Figure S4.** Structural verification with Ramachandran plots of EZH2-WT and modeled mutants V621M, A677M, R679C and H689Y after 50 ns of MD simulations (A), Ramachandran plot statistics percentage of residues plotted in most favored, additional allowed, generously allowed and disallowed regions of the plot.

**Figure S5.** Heatmap representation of RMSD fluctuations WT, MT structures and complexed with H3K27me0 peptide.

**Figure S6.** Secondary structure changes in EZH2-WT, EZH2-V621M, EZH2-A677T, EZH2-R679C and EZH2-H689Y when bound with H3K27me0 during 50 ns time period

**Figure S7.** Plot of first twenty eigen vector index and corresponding eigen values (A), projection of motion of the protein in the phase of first two principal eigen vectors of EZH2-WT, EZH2-V621M, EZH2-A677T, EZH2-R679C and EZH2-H689Y (B).

**Table S1.** Prediction of deleterious effects of 342 ns SNPs by PROVEAN server.

**Table S2.** Prediction score found to be significant by Polyphen 2 under HDivPred predictions

**Table S3.** Prediction score found to be significant by Polyphen 2 under HVarPred predictions.

**Table S4.** Protein BLAST search analysis of significant mutant sequences.
